# Systematic detection of abnormal samples reveals widespread mislabeling in metagenomic studies

**DOI:** 10.64898/2026.03.22.713545

**Authors:** Yong Zhou, Jun Chen, Wanxin Li, ShanShan Du, WeiMin Ye

## Abstract

The human microbiome plays a critical role in health and disease, and its dynamic nature has made longitudinal sampling a key strategy for elucidating microbiome–disease relationships. Although the gut microbiome generally stabilizes over time, a subset of samples frequently shows marked deviations from an individual’s baseline profile. We refer to these as *abnormal samples*. To analyze these abnormal samples, we developed a three-stage workflow to identify and classify these abnormal samples to figure out the underlying causes of these abnormal samples. Moreover, we systematically investigated abnormal samples across 16 publicly available metagenomic datasets, comprising a total of 5,171 metagenomes. Our analysis revealed that abnormal samples are often the result of mislabeling during sample collection, processing, or sequencing. Of which, fecal samples from family are more likely mislabeled. We found evidence of mislabeling in 75% of longitudinal datasets, involving up to dozens of samples per study, and in 25% of randomly selected cross-sectional datasets. Additional factors such as disease status (e.g., inflammatory bowel disease), sampling intervals, and sampling density may also contribute to sample abnormalities owing to true biological variations. These findings highlight that mislabeling is a common yet underrecognized issue in microbiome research. Our work underscores the importance of identifying and correcting abnormal samples to ensure data integrity in microbiome studies and provides a practical solution for quality control in large-scale metagenomic datasets.

## Introduction

The causal relationship between the human microbiome and disease has been extensively discussed in the literature^1–8^. This includes both direct gut-related diseases, such as IBS and IBD, and indirect conditions involving distant organs, like the brain or liver, through the gut-brain^9^ or gut-liver^10^ axis. Given the microbiome’s sensitivity to environmental exposures and its dynamic nature over time, many studies now employ longitudinal sampling to better characterize the association between microbiome composition and host phenotypes^3,5,7,8,11–13^. Generally, microbiome composition remains relatively stable within individuals over time, especially in adults.

However, abnormal samples are also observed in metagenomics studies, and these abnormal samples could be caused by environmental exposure such as antibiotic drugs or disease. In addition, stool samples often cannot be collected in a clinical setting. Participants are usually required to collect feces at home using a spoon-shaped tool, a process that can be uncomfortable and inconvenient. This makes consistent sampling challenging and increases the chance of cheating from participants, including the possibility of submitting another individual’s sample. Moreover, downstream steps—such as aliquoting, DNA extraction, library preparation, and sequencing—can introduce additional technical errors. Thus, the identification and further classification of abnormal samples and ruling out of the mislabels is essential for follow-up analyses.

Several strategies have been proposed to address this issue. One approach links host-derived DNA fragments in metagenomic data to matched host whole-genome sequencing (WGS) data^14^, achieving high sensitivity for sample identification. However, this strategy requires host WGS data, which is typically unavailable in most microbiome studies. Moreover, its specificity decreases among genetically related individuals, such as family members or individuals from homogeneous populations. Alternatively, host DNA can be extracted from stools and subjected to WGS for genotype comparison, but this method is often limited by the low abundance of human DNA in fecal samples. Without relying on host DNA, strain-tracking approaches have been used to identify shared microbial strains between samples^15–17^, The presence of identical strains suggests a common origin. However, strain-level comparison is computationally intensive, requiring pairwise genetic distance calculations across species, with time complexity scaling approximately as *O(M × N²)* (where *M* is the number of species and *N* the number of samples), making it impractical for large cohorts. In addition, many strain-tracking tools focus on dominant species or those with well-annotated reference genomes^18–20^, and they may be also confounded by inter-individual strain transmission, particularly among cohabiting or related individuals.

To address these limitations, we developed a three-stage workflow for systematic detection and classification of abnormal samples in metagenomic datasets. First, we introduced Find-abnormality, a distance-based, non-parametric tool that leverages ecological similarity structure to sensitively detect potential mislabels without requiring host genomic information. Second, we implemented a “place-back” strategy to distinguish true biological outliers from labeling errors. Third, we incorporated strain-level genetic distance analysis to further validate suspected mislabels.

Applying this workflow to both longitudinal and cross-sectional datasets, including 16 public metagenomic cohorts, we demonstrate that sample mislabeling is more prevalent than previously appreciated. Notably, samples from related individuals appear particularly susceptible to labeling errors. We further show that true biological abnormalities are associated with disease status, sampling interval, and temporal sampling density. Together, our study provides a practical framework for quality control in microbiome research and highlights the importance of rigorous abnormal-sample detection to ensure reproducible microbiome–disease associations.

## Results

### Identify and classify abnormal samples

The microbiome community profiles of abnormal samples often deviate substantially from baseline or expected patterns. To systematically detect such abnormalities, we developed a tool named *Find-abnormality*, which leverages Bray-Curtis dissimilarity to calculate pairwise distances between samples. For each sample, we ranked its dissimilarities to all other samples. Abnormal samples tend to show high distance rank, reflecting their dissimilarity from other samples of the same individual (**Figure 1A**). In contrast, normal samples from the same individual typically exhibit low distance rank. Briefly, for each individual with ≥2 samples, intra-individual Bray–Curtis distances were evaluated using a rank-based approach. For each pair of samples, the distance rank was calculated relative to all distances involving each sample. Sample pairs were considered inconsistent if their mutual distance exceeded the closest 5% of distances for both samples. Individuals containing at least one such inconsistent pair were flagged as potentially individuals. Within flagged individuals, graph-based clustering was performed by connecting highly similar sample pairs (mutual rank <10, **Figure 1A**). The largest connected component was considered the coherent longitudinal cluster, while remaining samples were classified as potential abnormal samples (**Figure 1A**)

**Figure 1.**
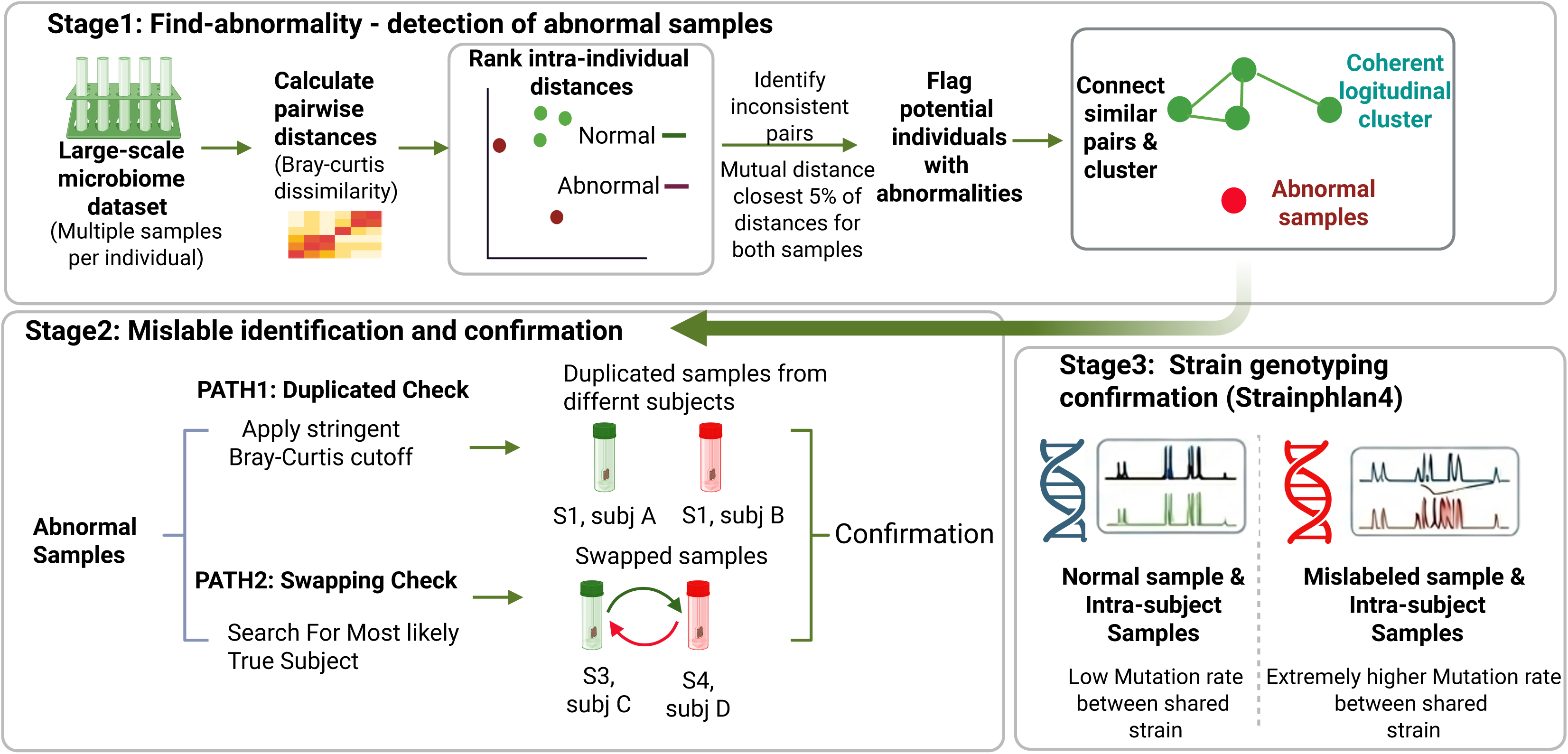
Overall Strategy for Systematic Microbiome Abnormality and Mislabel Detection. A three-stage workflow designed to identify and classify abnormal samples.(A) **Stage 1**: Detection of Abnormal Samples using Find-abnormality. Pairwise Bray-Curtis dissimilarities are calculated for all samples within a dataset. For each individual with multiple samples, the intra-individual distances are ranked relative to all distances involving those samples. Pairs are flagged as “inconsistent” if their mutual distance is high for both samples (above the 95th percentile). Within flagged individuals, graph-based clustering is used to identify a “coherent longitudinal cluster” (green nodes). Remaining unconnected samples are classified as “potential abnormal samples” (red nodes). (B) **Stage 2**: Mislabel Identification and Confirmation. Flagged potential abnormal samples from Stage 1 are further evaluated through two separate checks to identify mislabels. The Duplication Check (Path 1) uses a stringent Bray-Curtis dissimilarity cutoff to identify samples from different subjects that are highly similar, suggesting participant misconduct. The Swapping Check (Path 2) searches for the most likely true subject by looking for samples from other individuals that have a low distance rank to the abnormal sample. These identification steps are confirmed when re-assigning the sample leads to a normal intra-individual distance rank pattern. (C) **Stage 3**: Strain Genotyping Confirmation. The strain-level genotype of potential abnormal samples is compared using StrainPhlAn4. Shared strains between authentic intra-subject samples show a low mutation rate. In contrast, comparing a mislabeled sample to authentic intra-subject samples of its supposedly ‘true’ subject reveals an extremely higher mutation rate between shared strains, confirming the mislabeling.

Mislabels is one of abnormal samples that we need to focus on before the beginning of advanced microbiome studies. There exist two potential patterns for mislabel. One is the duplicated samples from different subjects, which may result from participant misconduct (e.g., submitting the same sample multiple times from different participants, **Figure 1B**). Another is the samples from different subjects are unintentionally swapped (**Figure 1B**). This is particularly relevant for samples taken at the same visit or baseline timepoint. For duplication check, we define a stringent Bray-cutis cutoff to identify the duplicated samples. Briefly, we sequenced dozens of same samples with different libraries and sequencing platforms, and compared distance between inter-subject, intra-subject and same-sample metagenomes to get a conservative cutoff to define the duplicated samples (**Method, Supplementary Figure 2**). For swapping check, we search for the abnormal samples’ most likely true subject based on distance rank value across different timepoints or batches, which is likely to happen from experimental errors during aliquoting. The abnormal samples are confirmed to be mislabel if the distance ranks among any paired samples within the swapped subjects become normal.

Finally, we further compared strain genotypes of abnormal samples using StrainPhlan4 (**Figure 1C**). The mutation rate of shared strains between samples is used to classify and confirm the mislabels (**Method**). The mislabeled sample and its intra-subjects’ samples tend to have extremely higher mutation rate (>0.1 mutation/kb) of species-level genome bins (SGBs) compared to normal ones (close to 0 mutation/kb). By employing this strategy, we could identify the abnormal samples in large-scale datasets and then narrow them to abnormal subjects and identify the mislabels. The overall design of this study is presented in **Supplementary Figure 1**, and a summary of the public datasets and associated metadata used is provided in **Supplementary Table 1**.

### Performance evaluation using simulated mislabeling in a longitudinal cohort

As mislabeling is an important source of abnormal samples and may significantly confound downstream analyses, we sought to determine whether the *Find-abnormality* approach in Stage 1 has sufficient power to recall abnormal samples caused by mislabeling. We used public dataset^11^ (PRJEB38984) as a test, which comprises gut metagenomes from a healthy Swedish population, as a benchmark cohort. Eighty individuals with four longitudinal samples were selected. Because samples were collected sequentially by visit, we simulated mislabeling at the visit (batch) level rather than across all samples simultaneously. The dataset consists of four batches—baseline, first visit, second visit, and third visit—with 80 samples per batch (**Figure 2A**). Strain-level SNP comparisons confirmed that all four time points originated from the same individual for each of the 80 subjects (**Methods**), ensuring a reliable ground truth.

**Figure 2.**
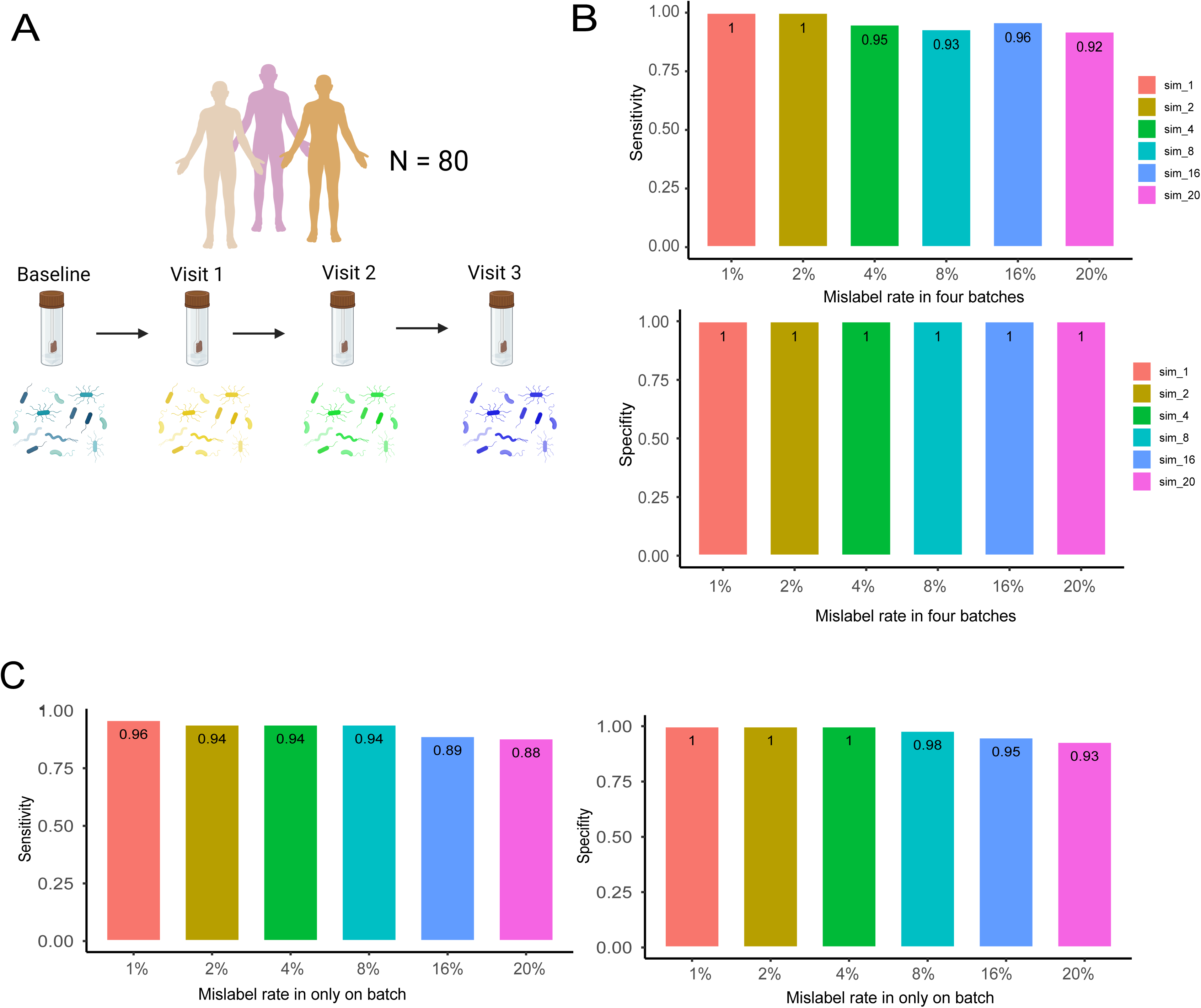
Performance evaluation of mislabel detection using simulated longitudinal data. (A) Study design for simulation analysis. Eighty individuals from the PRJEB38984 longitudinal cohort were selected, each with four time points (baseline, visit 1, visit 2, visit 3). Samples were organized into four visit-based batches (n = 80 per batch). Mislabeling was simulated at the batch level to mimic realistic experimental scenarios in which labeling errors occur during specific collection or processing stages. (B) Performance under simulated mislabeling across all batches. Mislabeling was introduced at varying rates (1%–20%) in all four batches simultaneously. Sensitivity (top) and specificity (bottom) are shown for each simulated error rate. The method maintained high performance across error levels, with specificity remaining 100% at lower mislabel rates and modest reductions observed as the mislabel proportion increased. (C) Performance under simulated mislabeling in a single batch. Mislabeling was introduced only within one visit batch at rates ranging from 1% to 20%. Sensitivity (left) and specificity (right) remained high across scenarios, achieving near-perfect performance at low mislabel rates (≤2%).

First, we simulated mislabeling in a single batch, reflecting a common scenario in microbiome studies. Under varying mislabel rates (1%–20%), the method achieved high sensitivity (92%–100%) and perfect specificity (100%, **Figure 2B**). Notably, when the mislabel rate was ≤2%, both sensitivity and specificity reached 100%. Second, we simulated mislabeling across all batches at proportions ranging from 1% to 20%. Compared to the single-batch scenario, performance showed a modest decline, with sensitivity ranging from 88% to 96% and specificity from 93% to 100% (**Figure 2C**). Specificity remained 100% when the mislabel rate was ≤4%, but both sensitivity and specificity gradually decreased as the mislabel rate increased. This reduction may partly reflect scenarios in which all samples from a given subject were consistently swapped with another subject, reducing detectable discordance within longitudinal trajectories.

Overall, these simulations demonstrate that our tool identifies mislabeled samples with high sensitivity and specificity, particularly when the mislabel rate is low, which is consistent with typical rates observed in real-world longitudinal microbiome datasets.

### Mislabels are prevalent in longitudinal gut metagenomes

To investigate abnormal samples in longitudinal metagenomic studies, we downloaded five publicly available datasets (PRJEB38984, PRJEB70966/PRJEB43119, PRJEB72385, PRJCA016454, and PRJCA026382) containing longitudinal samples from different diseases and analyzed them using the same pipeline (**Methods**). These datasets comprise 2,003 samples from 425 individuals **(Supplementary Table 1**).

Among them, PRJEB38984 includes 384 gut metagenomes from 101 normal gut. Notably, six paired metagenomes (3.1%) from different individuals were identified as duplicated samples (**Supplementary Table 2; Figure 3A**). To validate this observation, we compared the Bray–Curtis distances of these six paired metagenomes with 16 paired metagenomes derived from the same sequencing library of same sample and 240 paired metagenomes from different individuals. The Bray–Curtis distances of the six suspicious pairs were comparable to those observed between true intra-sample replicates and were markedly lower than inter-subject comparisons (**Figure 3B**), confirming that these six pairs were likely mislabeled. Furthermore, we quantified the mutation rates of shared SGBs between mislabeled metagenomes and true intra-subject samples. For example, W0020_3 was identified as a duplicated sample of W0075_3. The mutation rates of shared genotypes between mislabeled W0020_3 and its corresponding intra-subject samples were approximately tenfold higher than those observed among true intra-subject comparisons (**Figure 3C**). In contrast, normal W0075_3 and its longitudinal intra-subject samples exhibited comparable and extremely low mutation rates (<0.3 mutations/kb; **Figure 3D**). Similar patterns were observed in the other five duplicated samples (**Supplementary Figure 3**).

**Figure 3.**
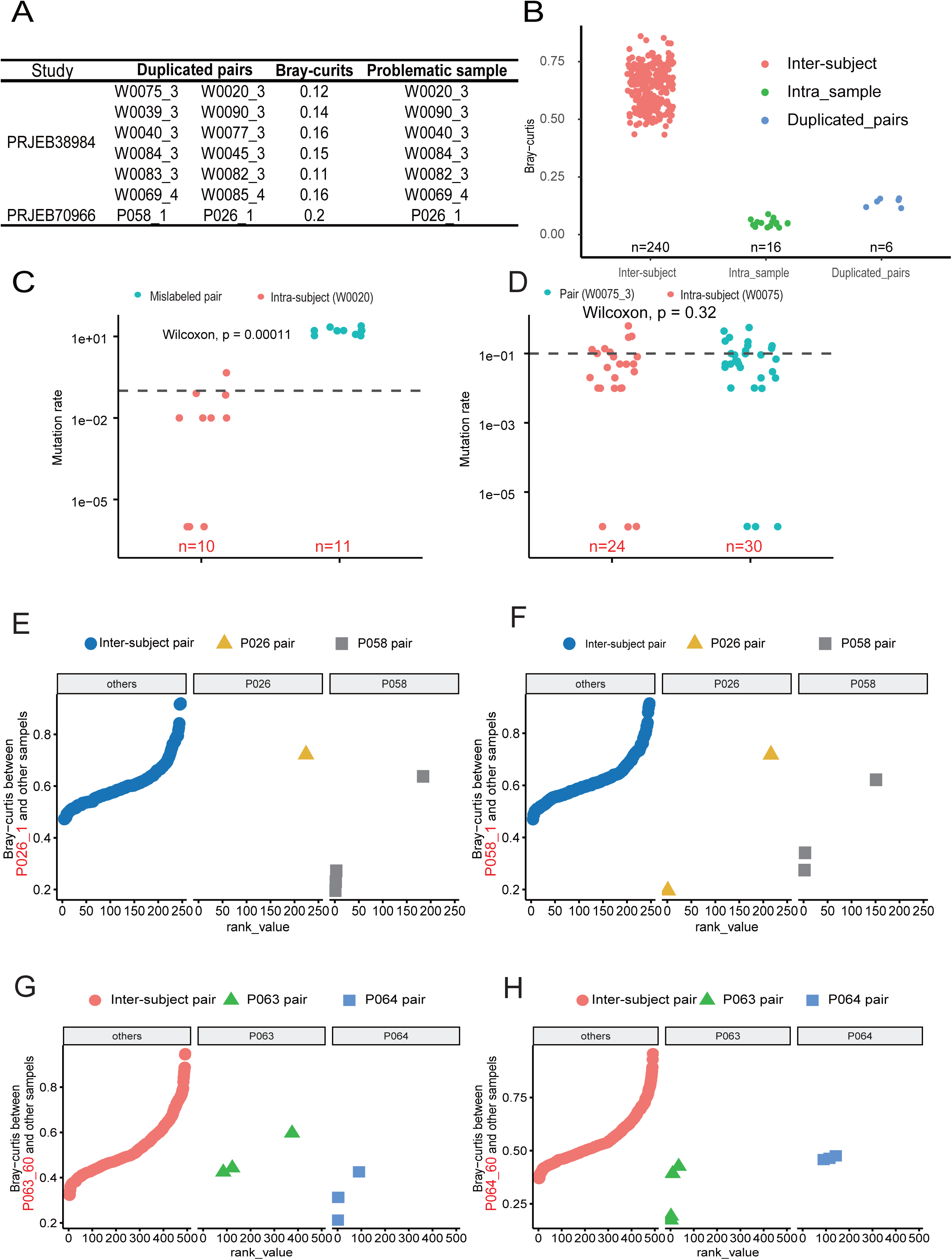
Identification and validation of mislabeled samples in longitudinal metagenomic datasets. (A) summary of duplicated samples across datasets. (B) Comparison of Bray–Curtis distances among three groups: true intra-individual longitudinal pairs (n = 16), inter-individual pairs (n = 240), and the six suspected duplicated inter-individual pairs (n = 6). (C) Mutation rates (log) of common strains between the mislabeled sample W0020_3 and its recorded intra-individual longitudinal samples, compared with mutation rates among true intra-individual longitudinal pairs. The dashed horizontal line indicates the predefined strain similarity cutoff of 0.1 mutations per kilobase. (D) Mutation rates (log) of shared strains among true longitudinal samples from the presumed source individual (W0075). (E). Bray–Curtis distance ranks of sample P026_1 relative to all inter-subject samples (“others”), its recorded intra-subject samples (P026), and samples from subject P058. (F) Bray–Curtis distance ranks for P058_1 show low ranks within its true intra-subject longitudinal samples. (G). Distance ranks of P063_60 relative to inter-subject samples (“others”), P063 samples, and P064 samples. (H). Distance ranks of P064_60 relative to inter-subject samples, P063 samples, and P064 samples.

PRJEB70966 comprises 248 longitudinal metagenomes collected before and after immune checkpoint therapy from 105 melanoma patients, with each individual contributing one to four time points. One of the 105 subjects (0.8%) was found to harbor a duplicated or mislabeled sample. Briefly, sample P026_1 was identified as likely originating from subject P058. The Bray–Curtis distance rank of P026_1 relative to its recorded intra-subject samples (P026) and to other inter-subject samples was high, indicating substantial compositional dissimilarity. In contrast, P026_1 showed a markedly low distance rank when compared to samples from subject P058 (**Figure 3E**), suggesting misassignment. Conversely, P058_1 showed low Bray–Curtis distance ranks relative to its true intra-subject longitudinal samples (**Figure 3F**), consistent with expected within-host temporal stability. Furthermore, strain-level analysis revealed that mutation rates across most SGBs between P026_1 and P058_1 were close to zero, indicating near-identical strain genotypes (**Supplementary Figure 5A**). Together, these findings strongly support that P026_1 was misattributed and most likely derived from subject P058. In addition, seven samples (2.8%) exhibited elevated intra-individual compositional dissimilarity (**Supplementary Table 3, Supplementary Figure 4**). Strain-level mutation rate analysis further supported mislabeling in these cases (**Supplementary Figures 5B and 6**).

PRJEB72385^21^ comprises 493 longitudinal samples from 124 individuals who received either fecal microbiota transplantation (FMT) or placebo capsules in a 1:1 randomized design. No duplicated metagenomes were detected in this cohort. However, eight samples (1.62%) were suspected to be mislabeled (**Supplementary Table 3**). Among these, P063_60 and P064_60 appear to have been swapped. Specifically, P063_60 exhibits a low Bray–Curtis distance rank relative to samples from individual P064 but a high distance rank relative to samples from P063 (**Figure 3G**). Conversely, P064_60 shows a low distance rank to samples from P063 and a high distance rank to samples from P064 (**Figure 3H**). These reciprocal patterns strongly support a sample identity swap between the two individuals. We also analyzed the mutation rate of SGBs between these eight samples and their corresponding intra-subject samples. One sample displayed same strains with its recorded intra-subject samples (**Supplementary Figure 7A**), suggesting biological continuity. In contrast, the remaining seven samples showed no evidence of same strains with their intra-subject samples (**Supplementary Figure 7B**), supporting their classification as likely mislabels.

The PRJCA016454 dataset comprises 272 longitudinal metagenomes from 76 individuals receiving weekly administration of histamine-2 receptor antagonists (H2RAs), proton pump inhibitors (PPIs), or no treatment. In this cohort, three individuals were identified as harboring suspected mislabeled samples. The microbiome composition of these abnormal samples was more similar to samples from other subjects than to their recorded intra-subject samples, consistent with potential sample misassignment (**Supplementary Figure 8**). The PRJCA026382 dataset includes 606 daily-collected metagenomes from 19 overweight or normal-weight individuals. Within this cohort, three individuals were found to contain one to two suspected mislabeled samples (**Supplementary Figure 9**).

Across the six longitudinal datasets analyzed, the proportion of mislabeled samples ranged from 1.62% to 3.6%, indicating that sample mislabeling is a pervasive issue in longitudinal metagenomic studies. Although the fraction of mislabeled samples appears small at the sample level, even a single mislabeled time point can propagate substantial computational bias across all longitudinal samples from the affected individual, particularly in analyses of within-host dynamics, strain evolution, and temporal stability. When evaluated at the individual level, 4%–16% of subjects across datasets were affected by at least one mislabeled sample (**Supplementary Table 2**), highlighting the disproportionate impact of sample misidentification on longitudinal analyses.

### Mislabel detection in CuratedMetagenomicData longitudinal cohorts

To further investigate sample mislabeling in longitudinal microbiome studies, we downloaded species-level abundance matrices and associated phenotype metadata from the CuratedMetagenomicData database. Among the available cohorts, three datasets contain well-annotated longitudinal information: HallAB_2017, HMP_2019_ibdmdb, and HMP_2019_t2d.

The HallAB_2017 dataset includes 259 stool samples from 20 patients with inflammatory bowel disease (IBD) and 12 controls. Samples were collected approximately monthly for up to 12 months, with the first collection occurring after treatment initiation. Application of our tool identified duplicated samples between subjects P0402 and P0403. Specifically, the Bray–Curtis dissimilarity among intra-subject samples from P0402 was comparable to the dissimilarity between samples from P0402 and P0403 (**Figure 4A**), indicating likely mislabeling between these two individuals. In addition, five subjects were identified as harboring abnormal samples. Detailed examination of species composition revealed that these abnormal samples exhibited markedly distinct community structures relative to their longitudinal counterparts (**Figure 4B–D; Supplementary Figure 10**). However, no clear swapped counterparts were detected for these samples, preventing definitive classification as mislabels. Notably, three of the five abnormal samples were dominated by only two or three species, suggesting potential sampling or sequencing artifacts rather than biological shifts (**Figure 4B–D**).

**Figure 4.**
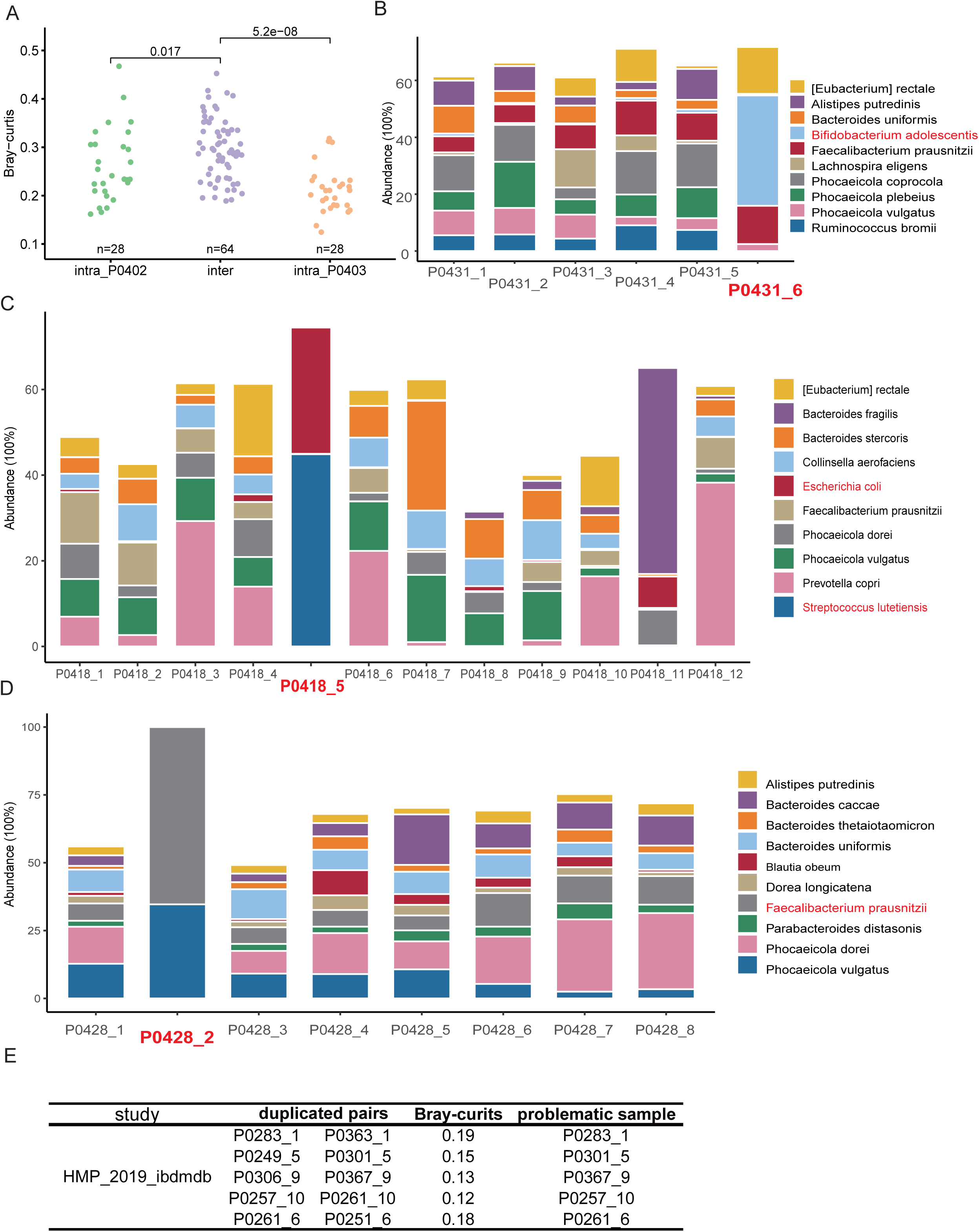
Identification of duplicated and abnormal samples in CuratedMetagenomicData longitudinal cohorts. (A) Detection of duplicated samples in the HallAB_2017 IBD cohort. Pairwise Bray– Curtis dissimilarities were calculated among intra-subject samples from P0402 (green), intra-subject samples from P0403 (orange), and inter-subject comparisons between P0402 and P0403 (purple). Statistical significance was assessed using two-sided tests (P values shown). (B–D) Representative abnormal samples identified in HallAB_2017. Stacked bar plots show relative species-level abundances across longitudinal samples from three individuals. Abnormal samples (highlighted in red) display marked compositional deviations compared with other time points from the same individual. (B) Example of a sample dominated by a limited number of taxa. (C) Example showing abrupt compositional restructuring relative to neighboring time points. (D) Example of extreme species predominance suggestive of potential sampling or sequencing artifacts. **(E)** Duplicated samples identified in the HMP_2019_ibdmdb cohort. The table summarizes duplicated subject pairs, corresponding Bray–Curtis dissimilarities, and the inferred problematic sample within each pair. Low dissimilarity values between different individuals support likely mislabeling events.

The HMP_2019_ibdmdb dataset comprises 1,585 samples from 130 individuals with IBD. Approximately 0.5% of subjects were found to harbor duplicated samples (**Figure 4E**). In total, 24 samples (1.3%) were flagged as abnormal. Among these abnormal samples, 12 exhibited extremely high compositional similarity to samples from other individuals (**Supplementary Figure 11**), strongly supporting the presence of true sample mislabeling events.

The HMP_2019_t2d cohort includes 189 longitudinal samples from 28 individuals with type 2 diabetes (T2D). No duplicated subjects were detected in this dataset. However, three samples (1.6%) from two individuals were identified as mislabeled: P0384_8, P0397_1, and P0397_11.

Compositional similarity analysis indicated that P0384_8 and P0397_1 are most consistent with subject P0379 (**Supplementary Figure 12A, B)**, whereas P0397_11 is highly similar to samples from subject P0395 (**Supplementary Figure 12C**), indicating likely cross-subject misassignment.

### Abnormal samples associated with disease

Abnormal samples may reflect true biological variation rather than sample mislabeling. In particular, specific disease states—such as chronic or acute infection, inflammatory conditions, or cancer treatment—can induce substantial shifts in gut microbial community composition. To investigate which diseases are more frequently associated with abnormal samples, we analyzed six longitudinal datasets with at least four time points per individual: four disease cohorts (IBD: HallAB_2017 and HMP_2019_ibdmdb; T2DM: HMP_2019_t2d and PRJCA026382), one healthy cohort (PRJEB38984), and one fecal microbiota transplantation (FMT) cohort.

Using Find-abnormality, we classified abnormal samples according to their temporal patterns. Abnormalities were categorized as either transient or persistent. We defined two or more consecutive abnormal time points within the same individual as representing true biological variation rather than isolated technical artifacts. In the two IBD datasets, consecutive abnormal samples were identified in 5/130 and 2/20 individuals, respectively. Among these eight individuals, seven exhibited persistent compositional shifts (**Figure 5, Supplementary Figure 13**), characterized by a marked change in microbiome structure followed by stabilization at a new community state. In contrast, two individuals displayed transient deviations, in which the microbiome underwent a substantial departure from baseline but subsequently returned to its original or stable configuration (**Figure 5A**, **Supplementary Figure 13**).

**Figure 5.**
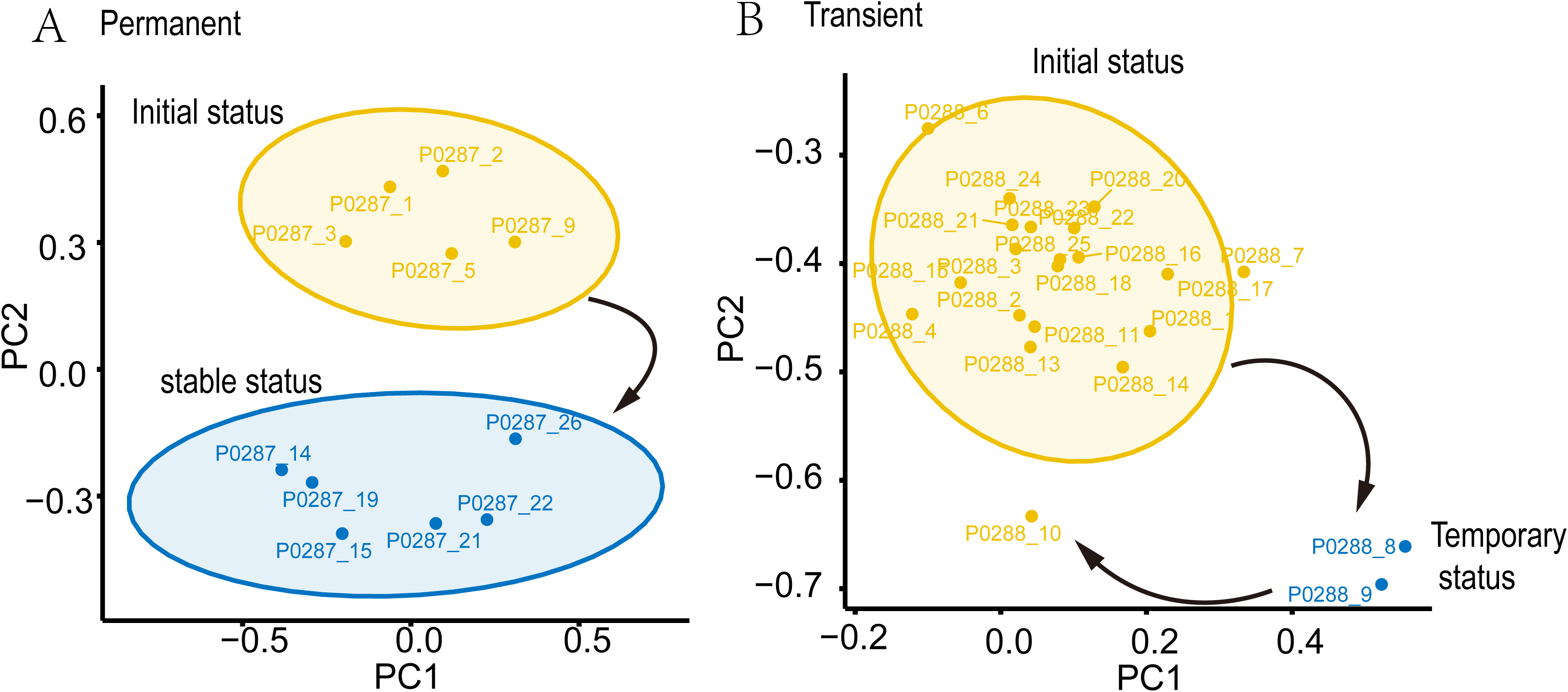
Temporal patterns of abnormal microbiome dynamics in longitudinal IBD datasets. Principal coordinate analysis (PCoA) of Bray–Curtis distances illustrating two distinct temporal patterns of abnormal samples identified by Find-abnormality. (A) Persistent shift. In some individuals, consecutive abnormal samples represent a sustained transition to a new microbiome configuration. The microbial community initially clusters within the baseline state (yellow ellipse, “initial status”) and subsequently shifts to a distinct, stable configuration (blue ellipse, “stable status”), indicating a long-term compositional change. (B) Transient deviation. In other individuals, abnormal samples represent temporary departures from the baseline microbiome state. Most time points cluster within the initial state (yellow ellipse), while a small number of samples deviate from this configuration (blue points, “temporary status”) before the microbiome returns to the baseline state. Sample labels correspond to individual identifiers followed by the sampling time order (e.g., _1, _2, _3 denote sequential time points for the same subject).

Notably, no persistent consecutive abnormalities were observed in the T2DM cohorts (HMP_2019_t2d and PRJCA026382) or in the healthy cohort (PRJEB38984). These results suggest that the gut microbiome remains relatively stable in healthy individuals and in T2DM cohorts under the studied conditions, whereas inflammatory diseases such as IBD are more likely to be associated with pronounced and sustained microbial community restructuring.

### Time interval and sampling density associated with abnormal samples

To investigate whether the time interval between consecutive sampling points influences the detection of abnormal samples, we evaluated microbiome community changes across different temporal intervals using an in-house longitudinal cohort. This dataset comprised 630 samples from 90 individuals in a natural population in Fujian Province (**Figure 6A**). Each individual contributed seven samples: five collected seasonally within one year and two collected annually in subsequent years (**Figure 6A**).

**Figure 6.**
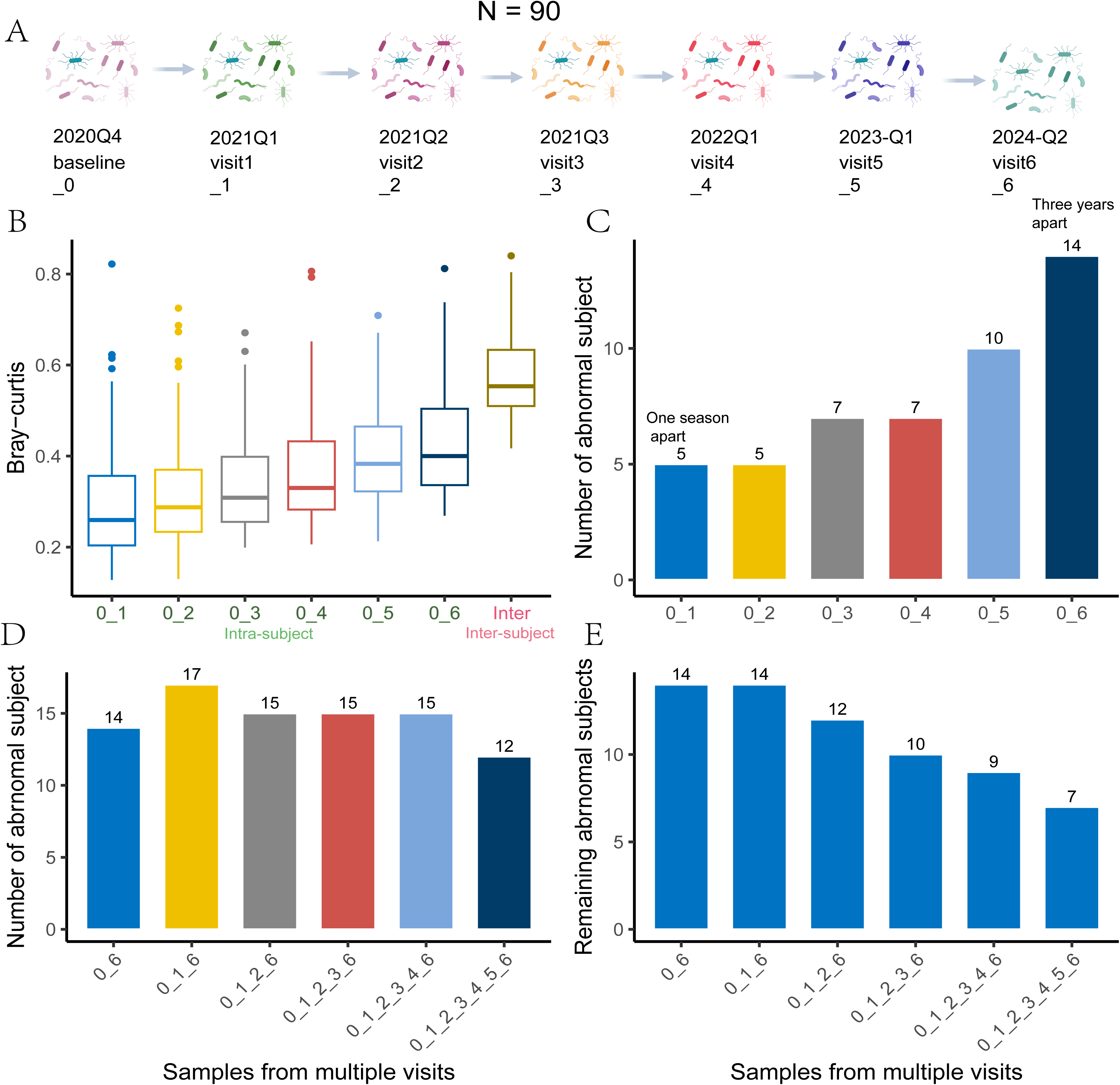
Time interval and sampling density influence abnormal sample detection in longitudinal microbiome analysis. (A) Study design of the Fujian natural population cohort (N = 90). Each subject contributed seven fecal samples: five collected seasonally within one year (2020Q4–2022Q1) and two additional samples collected annually thereafter (2023Q1 and 2024Q2). (B) Bray–Curtis distances between intra-individual sample pairs across increasing time intervals (0–1 through 0–6) compared with inter-individual distances (“inter”). Microbiome dissimilarity increases with time interval and gradually approaches inter-individual distances. Boxes represent interquartile ranges (IQR), center lines indicate medians, and whiskers denote 1.5 × IQR. (C) Number of subjects classified as abnormal when comparing pairs of samples separated by increasing time intervals. The frequency of abnormal classification increases with longer intervals, with >3-year intervals (0–6) showing nearly twice as many abnormal subjects as adjacent seasonal samples (0–1). (D) Number of abnormal subjects identified under different sampling-density scenarios, constructed by progressively including additional time points (e.g., 0–6; 0,1–6; 0,1,2–6; etc.). The overall number of abnormal subjects slightly decreases as sampling density increases. (E) Remaining abnormal subjects after stepwise inclusion of intermediate time points among the 14 initially identified abnormal subjects. Increasing sampling density reduces false abnormal classifications, indicating that denser longitudinal sampling improves stability of individual microbiome trajectory assessment.

As expected, microbiome dissimilarity increased with time interval. Intra-individual Bray–Curtis distances gradually approached inter-individual distances as the sampling interval lengthened (**Figure 6B**). Consistent with this trend, the number of samples flagged as abnormal increased with increasing time interval. Notably, pairs of samples separated by more than three years exhibited nearly twice as many abnormal classifications compared to pairs collected one season apart (**Figure 6C**), suggesting that longer temporal gaps increase the likelihood of apparent anomalies.

To further examine the impact of sampling density on abnormal sample identification, we conducted stepwise analyses by progressively increasing the number of time points per individual. Although the total number of subjects classified as abnormal slightly decreased overall (**Figure 6D**), we observed that 14 individuals initially flagged as abnormal were reclassified as normal when additional intermediate time points were included (**Figure 6E**). Collectively, these results demonstrate that both time interval and sampling density substantially influence abnormal sample detection in longitudinal microbiome studies.

### Application in datasets with cross-sectional gut metagenome

Although primarily designed for longitudinal analyses, our tool can also identify duplicated and mislabeled samples in cross-sectional metagenomic datasets. To evaluate its broader applicability, we randomly selected eight publicly available gut microbiome projects from three major repositories (EBI, NCBI, and GSA): PRJNA613947^15^, PRJNA401977^22^, PRJNA1065729, PRJNA397906, PRJEB18755, PRJCA004518, PRJCA005417, PRJCA009886. These studies comprise 927 metagenomic samples (**Supplementary Table 4**). and were originally designed as case–control comparisons to identify disease-associated microbial species or pathways.

Our analysis detected duplicated samples in two datasets. In PRJNA613947, 12 metagenomes (3.5%,6 pairs) were identified as duplicates, while PRJNA401977 contained 4 duplicated metagenomes (1.1%, 2 pairs) (**Figure 7A**). Strain-level analysis showed that shared strains between duplicated pairs exhibited extremely low mutation rates (<1 mutation per kilobase), consistent with a common biological origin (**Figure 7B–C**). In PRJNA613947, a study comparing gestational diabetes mellitus (GDM) with non-GDM controls, one GDM sample was mislabeled as a non-GDM control (**Supplementary Table 5**). Additionally, within dataset PRJNA613947 of longevous individuals and their offspring in the same project, three of six duplicated pairs originated from members of the same family (**Figure 7C**), suggesting that samples from related individuals may be particularly susceptible to labeling errors. Collectively, these findings demonstrate that duplicated and mislabeled samples are present in publicly available cross-sectional metagenomic datasets and may confound case–control analyses if not properly detected and corrected.

**Figure 7.**
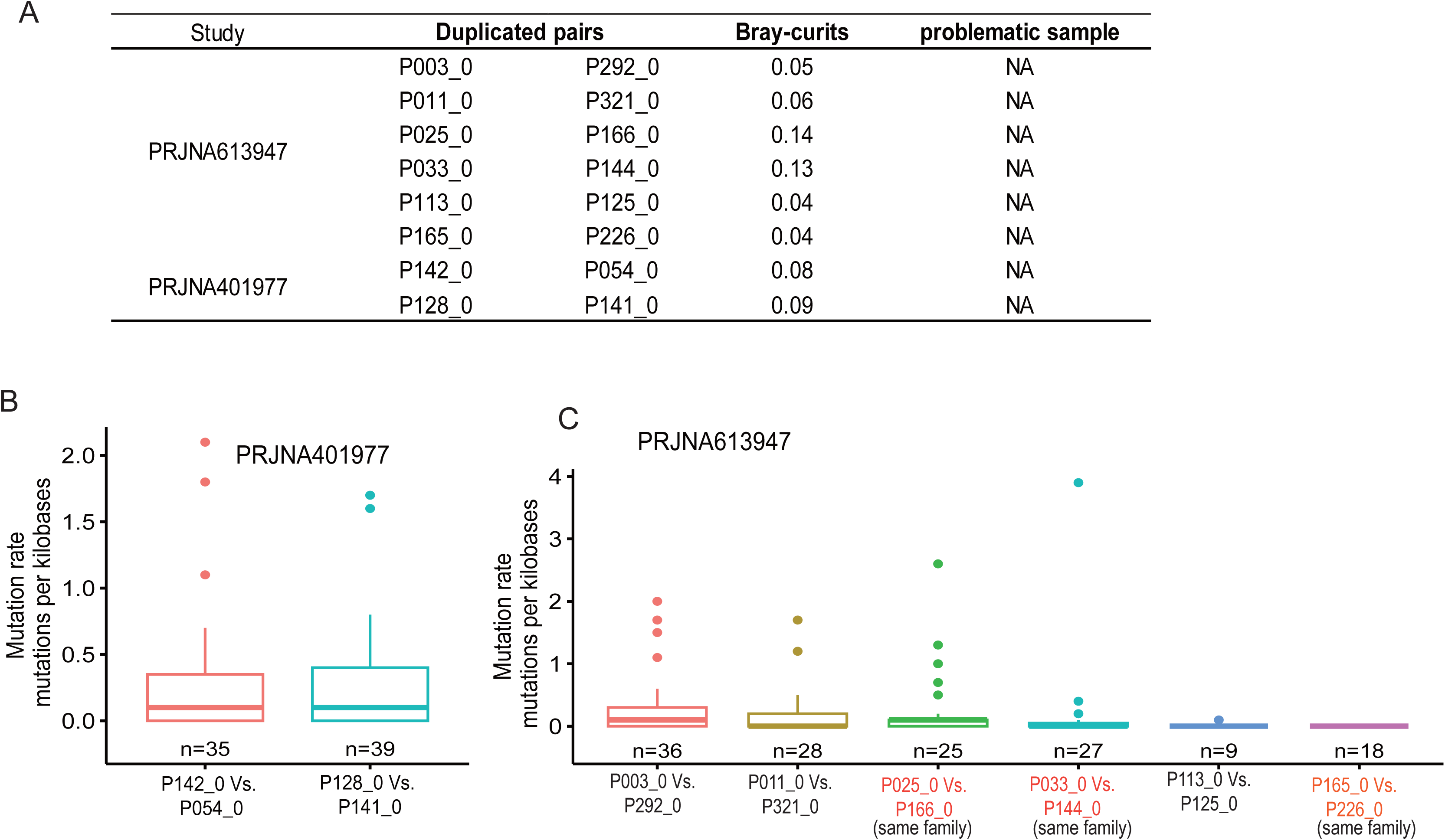
Identification of duplicated and mislabeled samples in public cross-sectional metagenomic datasets. (A) Summary of duplicated sample pairs detected in PRJNA613947 and PRJNA401977. For each pair, the Bray–Curtis distance is shown. All duplicated pairs exhibit extremely low community dissimilarity, consistent with potential sample duplication. (B) Strain-level mutation rates between duplicated samples in PRJNA401977. Mutation rates are calculated as the number of single-nucleotide variants per kilobase in shared strains. Duplicated pairs display very low genomic divergence (<1 mutation per kilobase), supporting a common biological origin. Boxplots show median and interquartile range; whiskers indicate 1.5 × IQR; points represent individual strains. (C) Strain-level mutation rates for duplicated pairs in PRJNA613947. Most duplicated pairs show minimal genetic divergence. Three duplicated pairs derived from members of the same family are highlighted. Sample sizes (number of shared strains) are indicated below each comparison.

## Discussion

Abnormal samples are frequently observed in longitudinal microbiome studies. Identifying mislabeled samples among these abnormalities can significantly enhance our understanding of intra-individual heterogeneity in microbiome communities. In this study, we developed a workflow to systematically investigate abnormal samples across 16 microbiome datasets. Our findings indicate that most abnormalities can be attributed to mislabeling during sample collection or genome sequencing. Notably, a substantial portion of these mislabels appear to be duplicated samples, suggesting potential intentional duplication or misconduct by sample donors. This is supported by the frequent detection of duplicated samples among family members in dataset PRJNA613947. In addition to technical artifacts, true biological variation—driven by disease progression, sampling frequency, and time intervals—may also contribute to abnormal samples. These results emphasize the value of analyzing, rather than excluding, abnormal samples, as they can reflect either critical biological variation or procedural errors.

Several limitations of our tool and workflow should be acknowledged. First, validation of mislabels in the later stages relies on strain tracking, which assumes that samples originating from the same individual share at least a subset of strain genotypes (≥2 shared strains). This assumption enables differentiation between true biological variation and sample mislabeling. However, it may be violated in situations where unrelated samples coincidentally share identical strains due to horizontal transmission, particularly among family members or cohabiting individuals^23–25^. In such contexts, strain sharing may reduce the specificity of mislabel confirmation. Second, some abnormal samples identified in stage 1 could not be conclusively validated in stages 2 or 3. For instance, we did not observe reciprocal sample swaps within the same dataset, nor did we consistently detect shared strains between abnormal samples and their corresponding intra-individual samples. In certain cases, abnormal samples were characterized by markedly reduced species diversity and dominance of one or a few taxa, patterns that may indicate DNA extraction failure or other severe technical artifacts. Alternatively, these samples could result from mislabeling involving external samples not included in the analyzed dataset, or from genuine biological perturbations. Although such samples were not formally classified as mislabels in this study—so as to maintain conservative mislabeling estimates—we believe that a subset likely reflects technical failure or labeling errors. Finally, our workflow is predicated on the relative temporal stability of the adult gut microbiome and may not be directly applicable to infant microbiome studies. During early life, microbial communities are highly dynamic and less stable, which may increase the risk of false-positive mislabel classifications. Therefore, caution should be exercised when applying this framework to very young populations.

Controlling mislabels in gut microbiome studies is essential. First, stool sample collection is inherently difficult, particularly in large-scale nature population studies, where participant compliance may be limited. Second, the multi-step processes from sample collection to sequencing introduce multiple opportunities for mislabeling. Implementing automated workflows during sample preprocessing and sequencing could help reduce these errors. Third, even correctly labeled samples may not reflect the individual’s stable microbiome state, as microbiome composition can fluctuate significantly. Increasing sampling density in longitudinal studies may offer a more accurate characterization of stable microbial communities and improve identification of true biological variations instead of negative variations from mislabels.

## Methods

### Sample processing and metagenomic sequencing

A total of 90 individuals (aged 35–70 years) from Gaoshan Town, Fujian Province, China, were enrolled in this study. Fecal samples were collected and processed for metagenomic sequencing. Genomic DNA was extracted from 1 mL of fecal suspension using a two-step protocol. First, samples were incubated with a lysozyme lysis buffer (20 mg/mL lysozyme) at 37 °C for 60 min to facilitate enzymatic cell wall disruption. Mechanical lysis was subsequently performed using 1.0-mm diameter glass beads in a FastPrep 24-5G Homogenizer (MP Biomedicals). In the second step, DNA was purified using the MGIEasy II kit according to the manufacturer’s instructions. DNA concentration was quantified using a Qubit 4.0 Fluorometer (Life Technologies). Sequencing libraries were constructed using the DNA Nanoball (DNB, BGI) preparation protocol following the manufacturer’s guidelines. Libraries were sequenced on the MGISEQ-T7 platform according to standard protocols.

Sixteen samples were sequenced twice using the same DNA library, and sixty-two samples were sequenced twice using independently prepared DNA libraries to determine the cutoff for duplicated samples. The remaining samples were sequenced once using newly prepared DNA libraries. All study procedures were approved by the Biomedical Ethics Review Committee of Fujian Medical University (Approval No.: IACUC FJMU 2023-0058).

### Datasets collection

In this study, we have collected 8 datasets with longitudinal metagenomes consisting of 4307 samples. The abundance and metadata of the other three datasets, HallAB_2017, HMP_2019_idbmdb, and HMP_2019_t2d, are from the CuratedMetagenomicData database. Raw fastq data from the other 13 datasets are downloaded for EGA, NCBI or GSA database. In addition, we have also collected raw fastq data from eight datasets with cross-sectional samples only. These comprise 928 samples ranging from 39 to 348 for each dataset. The eight cross-sectional and five longitudinal datasets above have been reanalyzed using the same pipeline. Of note, for datasets with raw fastq data, a proportion of samples could not be downloaded correctly. These samples are ruled out for this study.

### Species and Strain-level profiling of metagenomics samples

Metagenomic data preprocessing was performed using KneaData^26^ with default parameters. Briefly, low-quality bases were trimmed from the ends of reads, and host-derived sequences were removed by aligning reads to the human reference genome (hg19) using Bowtie2². The resulting high-quality, host-depleted reads were retained for downstream analyses. Species-level taxonomic profiling was conducted using MetaPhlAn 4^27,28^ with default parameters and the SGB database (mpa_vOct22) based on clean FASTQ data.

Strain-level profiling was performed using StrainPhlAn4^27,28^ with the custom SGB marker database (mpa_vOct22) and the parameters “-mutation_rates marker_in_n_samples 1 -sample_with_n_markers 10 –phylophlan_mode accurate”. Because pairwise mutation rate estimates may vary slightly depending on the total number of samples included in the multiple-sequence alignment step of StrainPhlAn 4, we controlled for this effect by limiting the number of input samples per SGB (abundance > 0) to approximately 100 whenever possible.

### Identification of abnormal samples using Find-abnormality

We developed a Python-based quality control pipeline to detect mislabeled individuals and duplicate samples in longitudinal metagenomic datasets. The workflow integrates ecological distance metrics, rank-based anomaly detection, and graph-based clustering. Species-level abundance matrices (features × samples) were log-transformed to reduce compositional skew. Pairwise sample dissimilarities were computed using Bray–Curtis distance, generating a symmetric distance matrix.

For each individual with ≥2 samples, intra-individual Bray–Curtis distances were evaluated using a rank-based approach. For each pair of samples, the distance rank was calculated relative to all distances involving each sample. Sample pairs were considered inconsistent if their mutual distance exceeded the closest 5% of distances for both samples. Individuals containing at least one such inconsistent pair were flagged as potentially mislabeled. Within flagged individuals, graph-based clustering was performed by connecting highly similar sample pairs (mutual rank <10). The largest connected component was considered the coherent longitudinal cluster, while remaining samples were classified as potential mislabels.

Within each sequencing batch, sample pairs with Bray–Curtis distance below a predefined cutoff (default = 0.21) were flagged as potential duplicates. When multiple batches were available, the cutoff could alternatively be estimated as the mean of the five smallest intra-individual adjacent-sample distances from non-problematic individuals. For each putative mislabel, the mean distance rank to samples from every individual was calculated. Individuals were ranked by proximity, and the top candidates were reported as potential true labels.

### The cutoff of duplicated samples identification

To determine the Bray–Curtis distance cutoff for identifying duplicated samples, we performed repeated sequencing of a subset of samples using either the same DNA library (N = 16) or independently prepared libraries from the same DNA extract (N = 62). Bray–Curtis distances were calculated between these duplicated samples. As expected, duplicated samples sequenced from the same library exhibited significantly lower distances than those sequenced from different libraries.

The Bray–Curtis distances between duplicated samples ranged from 0.04 to 0.27 **(95% CI)** (**Supplementary Figure 2A**), which were markedly lower than distances observed between samples from different individuals (**95% CI: 0.50–0.78**). We further compared these values with intra-individual distances from longitudinal samples of 80 individuals described above. The duplicated-sample distances were clearly separated from inter-individual distances (95% CI: 0.43–0.68). To minimize false-positive detection, we defined the 90th percentile of the duplicated-sample distance distribution as the default Bray–Curtis cutoff for identifying duplicated samples (**Supplementary Figure 2B**).

### Leveraging strain mutation rate to confirm sample source

We assume that samples from the same individual have a lower mutation rate, which is significantly lower than that between samples of different individuals. The mutation rate between two samples is calculated as the mutation number per a thousand nucleotide bases obtained through Strainphaln4. In line with the assumption, we observed that the mutation rate for each SGB between two samples from the same individual is less than 0.1 mutation/kb (**Supplementary Table 6**). In contrast, the mutation rate of SGB for samples from different individuals is far beyond one mutation/kb. Using this cutoff, whether the samples from the same individual could be confirmed.

### Statistics

Python (v3.9) is used to preprocess the data, and R (4.3) is used to visualize the data. p-value is given and thought of as significant if p < 0.05. The two-group comparison uses The Wilcoxon Rank Sum Test in R.

## Code availability

The tool Find-abnormality and metadata and species profiles of 16 datasets are available in github: https://github.com/zhouyonog0713/Find-abnormality. The raw data is under GSA: https://ngdc.cncb.ac.cn/gsa/s/d63LF1Eq.

## Funds

This study was jointly supported by National Key R&D Program of China(2024YFC3405800, Weiming Ye); General Program of the Natural Science Foundation of Fujian Province [grant number: 2023Y9116 Weiming Ye; 2024J01577 Wanxin Li]; Government of Fuqing city [grant number: 2019B003 Weiming Ye], and High-level Talents Research Start-up Project of Fujian Medical University (No. XRCZX2020037 Wanxin Li XRCZX2022001 Jun Chen, XRCZX2023030 Weiming Ye, XCRZX2023004 Yong Zhou, and XRCZX2023005 Shanshan Du).

## Supplementary Figures

**Supplementary Figure 1.**
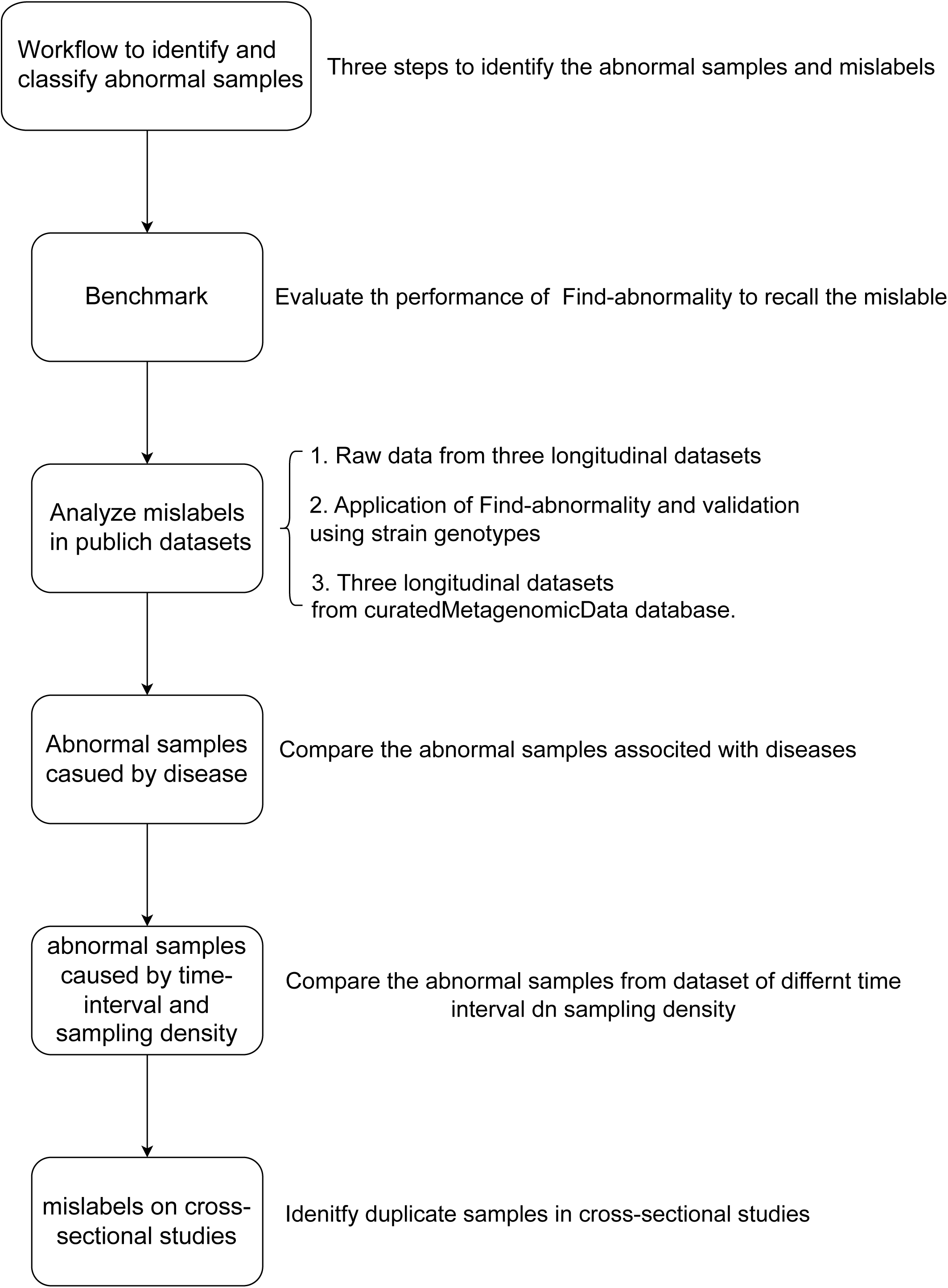
Overview of the study workflow for abnormal sample detection and validation. Flowchart illustrating the analytical framework used in this study. First, the workflow was developed to identify and classify abnormal samples based on ecological distance profiles. Second, a mislabel simulated strategy was implemented to evaluate performance by simulating random labeling errors at varying proportions. Third, the workflow was applied to public longitudinal datasets, including raw data from three independent cohorts and three additional datasets from the curatedMetagenomicData database, with mislabels validated using strain genotyping. Fourth, abnormal samples associated with disease status were evaluated by comparing datasets across different disease conditions. Fifth, the influence of sampling interval and sampling density on abnormal sample detection was assessed using longitudinal cohorts with varying temporal designs. Finally, the framework was extended to eight cross-sectional datasets, where potential mislabels were detected and validated using strain-level genetic similarity (StrainPhlAn4).

**Supplementary Figure 2.**
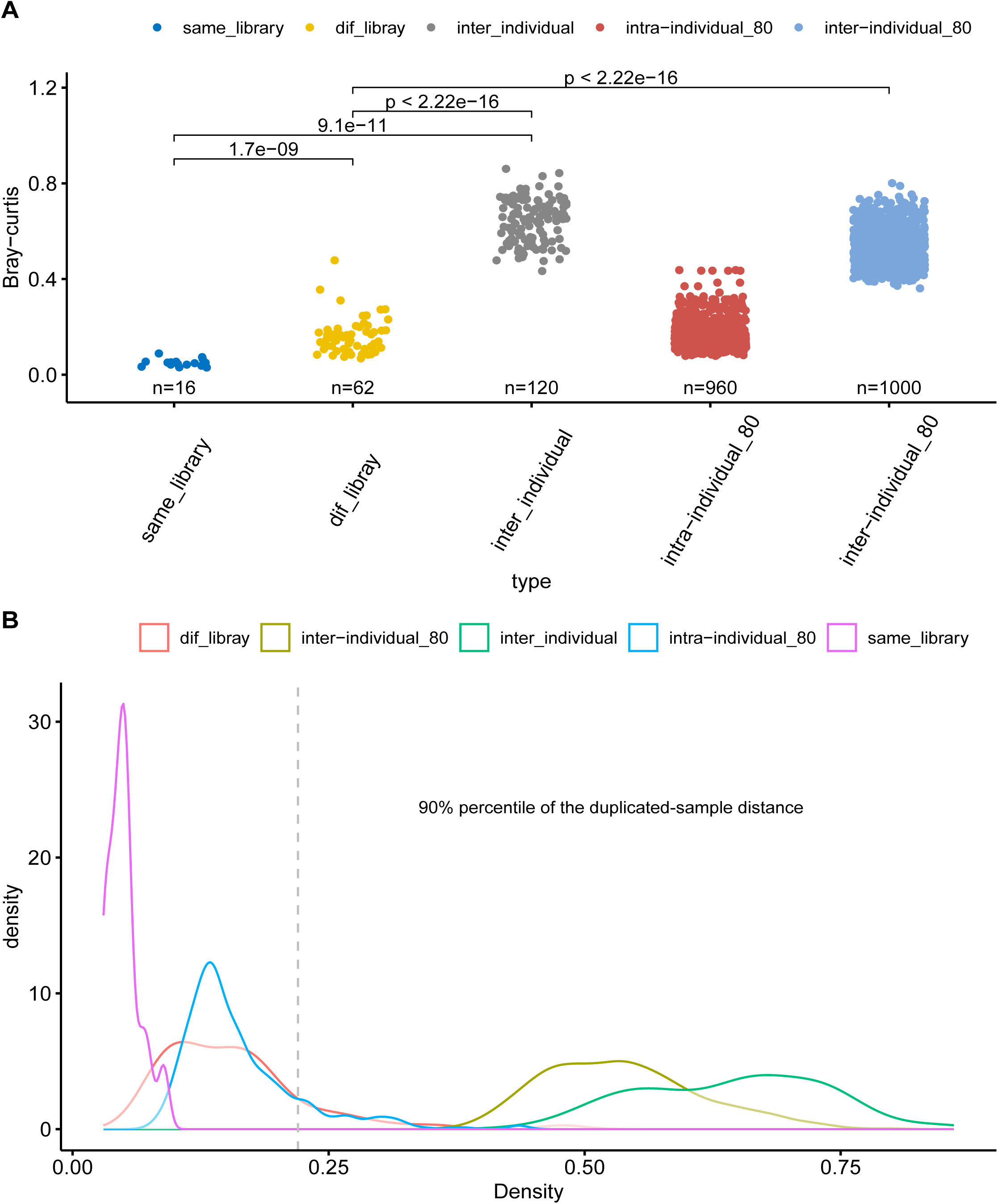
Determination of the Bray–Curtis cutoff for identifying duplicated samples. (A) Bray – Curtis distances among different comparison groups. Distances were calculated between samples sequenced twice using the same library (same_library), using independently prepared libraries from the same DNA extract (dif_library), between samples from different individuals (inter_individual), and between longitudinal samples from the same individual (intra-individual_80) or different individuals (inter-individual_80) in an independent cohort of 80 subjects. P-values were calculated using Wilcoxon rank-sum tests. Sample sizes are shown below each group. (B) Density distributions of Bray–Curtis distances for the same comparison groups. The vertical dashed line indicates the 90th percentile of the repeated-sample distance distribution, which was selected as the default cutoff for identifying duplicated samples. This threshold clearly separates duplicated samples from inter-individual distances while minimizing false-positive detections.

**Supplementary Figure 3.**
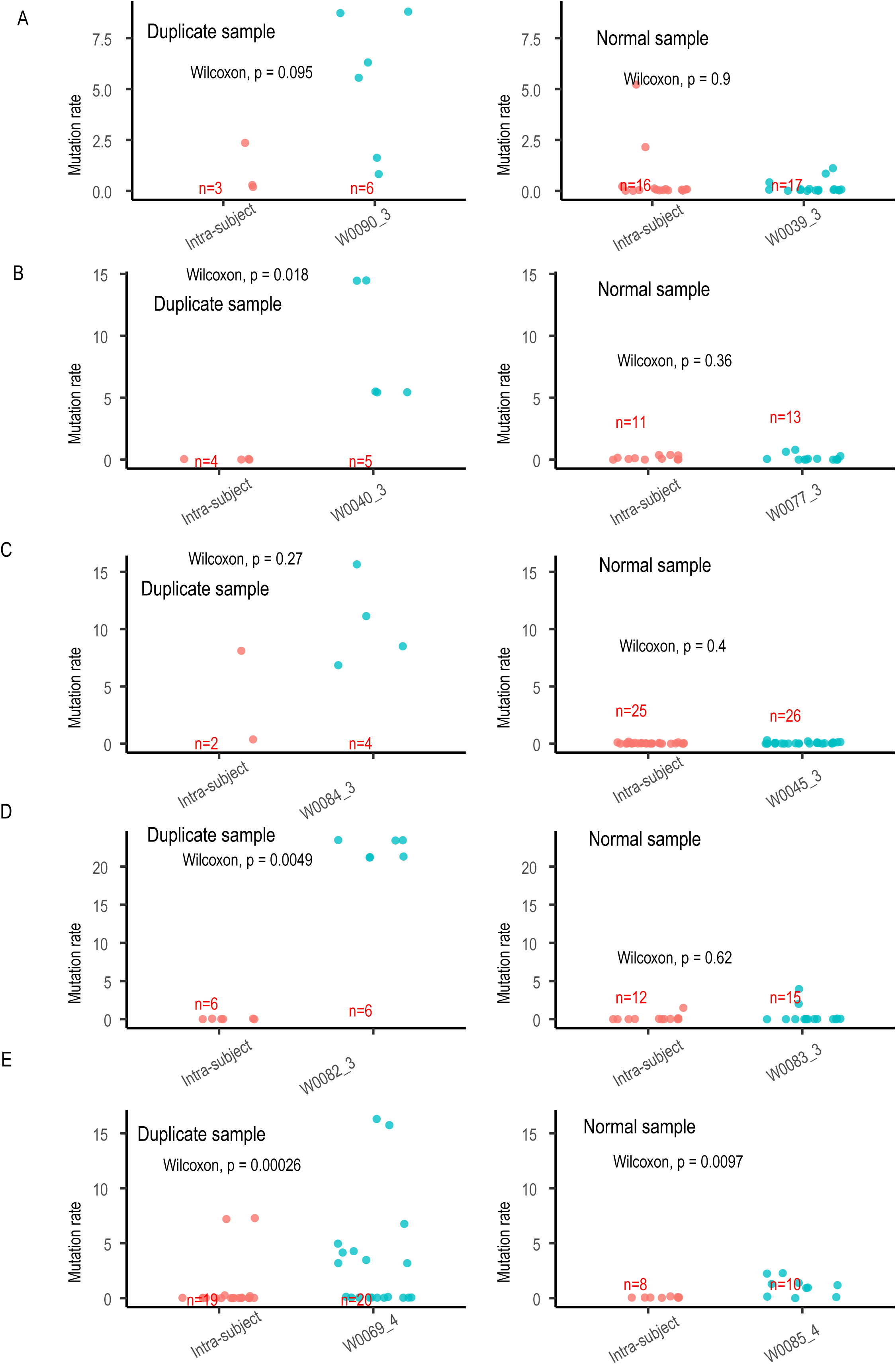
Strain-level mutation rate comparisons between suspected duplicated samples and intra-subject baseline samples. Strain-level mutation rates were calculated for common SGBs. Each row (A–E) presents one representative case. For each case, the left panel shows a suspected duplicated sample, and the right panel shows a confirmed normal sample as a control comparison. Mutation rates between the queried sample (e.g., W0000_3, W0040_3, W0084_3, W0082_3, W0069_4) and their intra-subject samples are shown in blue. Mutation rates between samples (not contain queried sample) from the same subject (intra-subject baseline) are shown in red. Wilcoxon rank-sum test p-values are indicated in each panel. Sample sizes (n) for each comparison group are shown below the x-axis.

**Supplementary Figure 4.**
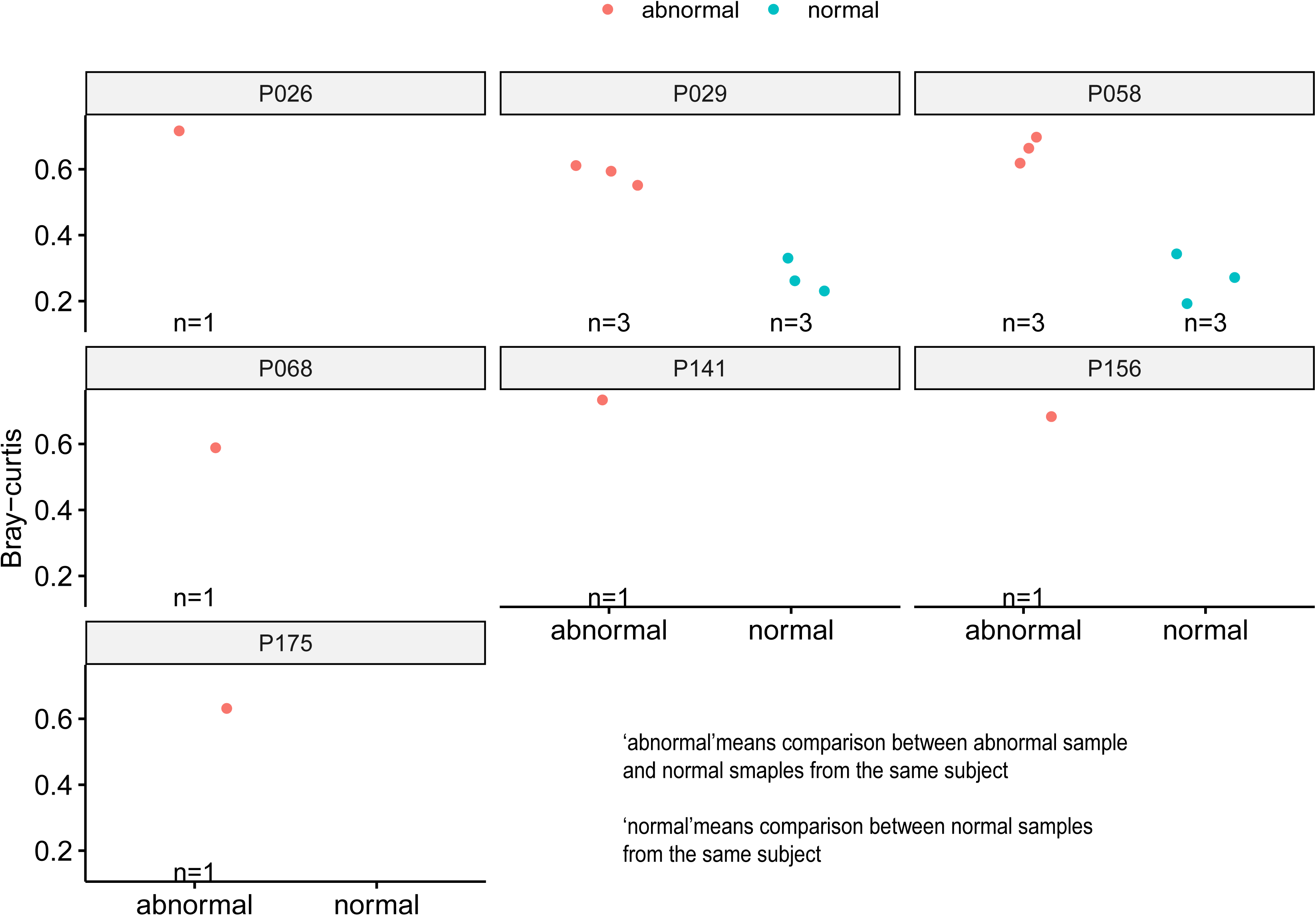
Elevated intra-individual Bray–Curtis dissimilarity in additional abnormal samples from PRJEB70966. Bray–Curtis distances within individual subjects from PRJEB70966 are shown for seven additional samples exhibiting unusually high intra-individual dissimilarity. Each panel represents one subject. Red points (“abnormal”) denote Bray–Curtis distances between the abnormal sample and the subject’s other longitudinal samples. Blue points (“normal”) represent Bray–Curtis distances among normal longitudinal samples from the same subject (intra-subject baseline). In some subjects, blue points are absent because only one normal sample was available, precluding intra-subject baseline comparisons. Sample sizes (n) for each comparison are indicated below the x-axis.

**Supplementary Figure 5.**
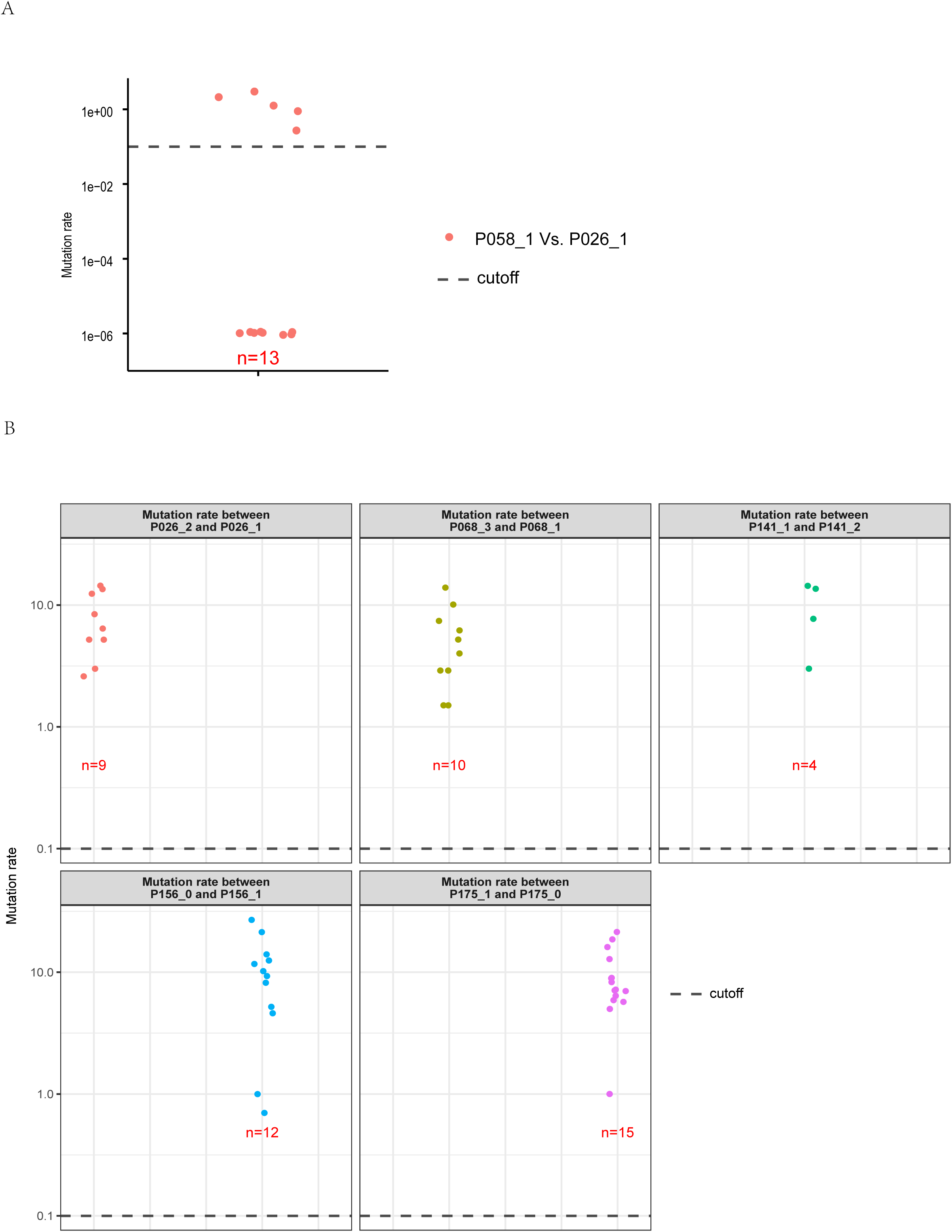
Strain-level mutation rate analysis supporting mislabel identification in PRJEB70966. (A) Mutation rates across shared SGBs between samples P058_1 and P026_1. Each point represents the mutation rate (log scale) for one SGB. The dashed horizontal line indicates the predefined strain similarity cutoff of 0.1 mutations per kilobase. Most shared SGBs exhibit mutation rates close to zero and below this cutoff, indicating near-identical strain genotypes between P058_1 and P026_1. The number of shared SGBs analyzed (n) is indicated. (B) Mutation rates between additional abnormal sample pairs identified in PRJEB70966 (P026_2 vs P026_1, P068_3 vs P068_1, P141_1 vs P141_2, P156_0 vs P156_1, and P175_1 vs P175_0). Each point represents one shared SGB. The dashed horizontal line denotes the same cutoff (0.1 mutations per kilobase). In contrast to panel A, most mutation rates exceed the cutoff, indicating distinct strain genotypes and supporting mislabeling or severe sample inconsistency rather than true longitudinal continuity. Sample sizes (n) for each comparison are shown in red.

**Supplementary Figure 6.**
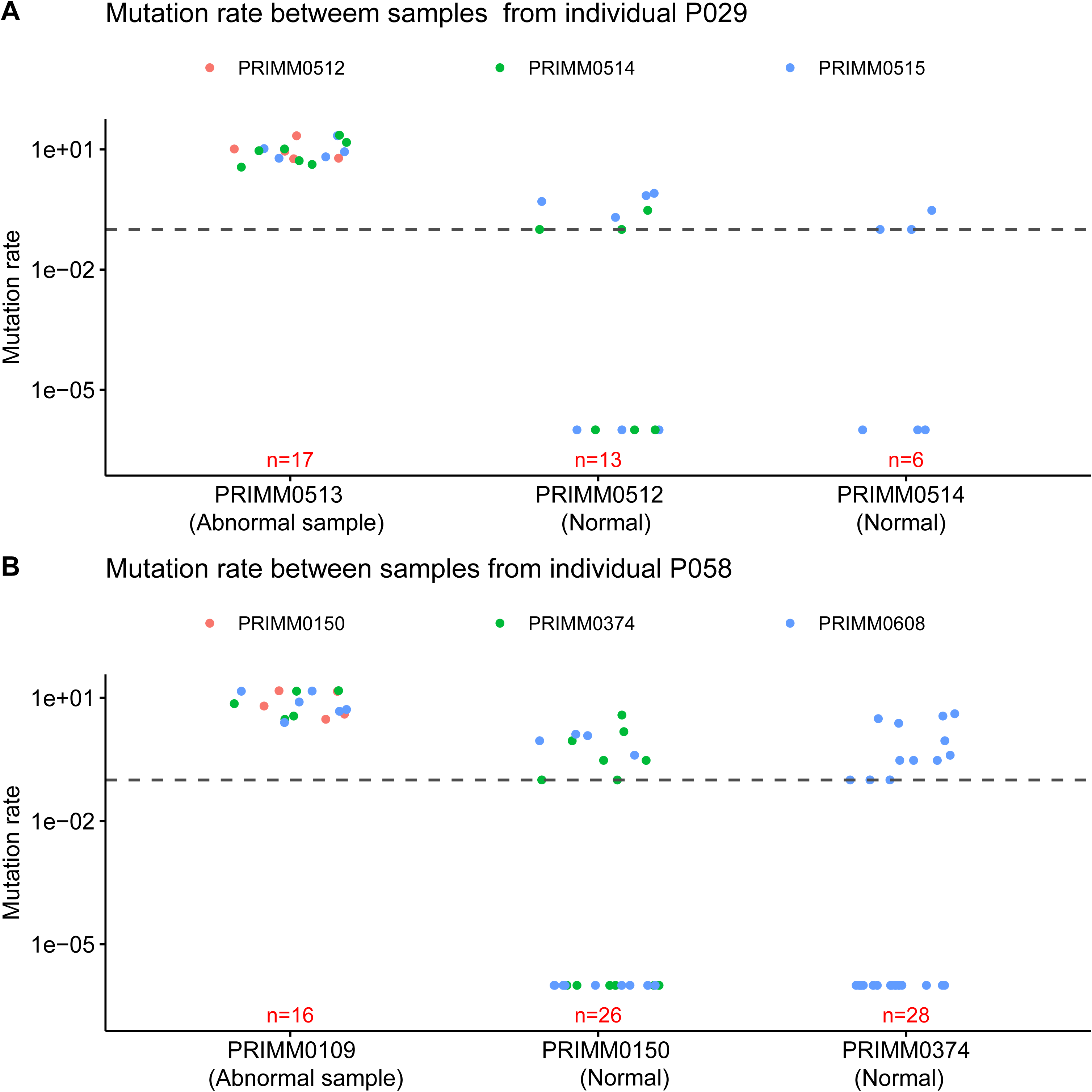
Mutation rates of strain genotypes for additional two subjects with abnormal samples. (A) Mutation rates (log) between samples from individual P029. PRIMM0513 (labeled abnormal) is compared against other samples from the same subject, showing mutation rates predominantly above the cutoff, consistent with strain-level divergence. In contrast, comparisons among normal samples (PRIMM0512 and PRIMM0514) exhibit mutation rates largely below the cutoff, indicating expected intra-subject strain continuity. Points are colored by the comparison partner as indicated in the legend. Sample sizes (number of shared SGBs) are shown below each comparison. (B) Mutation rates (log) between samples from individual P058. Similar to panel A, the abnormal sample PRIMM0109 shows elevated mutation rates relative to other samples from the same subject, whereas comparisons among normal samples (PRIMM0150 and PRIMM0374) display mutation rates largely below the cutoff. The dashed horizontal line represents the predefined strain similarity cutoff of 0.1 mutations per kilobase.

**Supplementary Figure 7.**
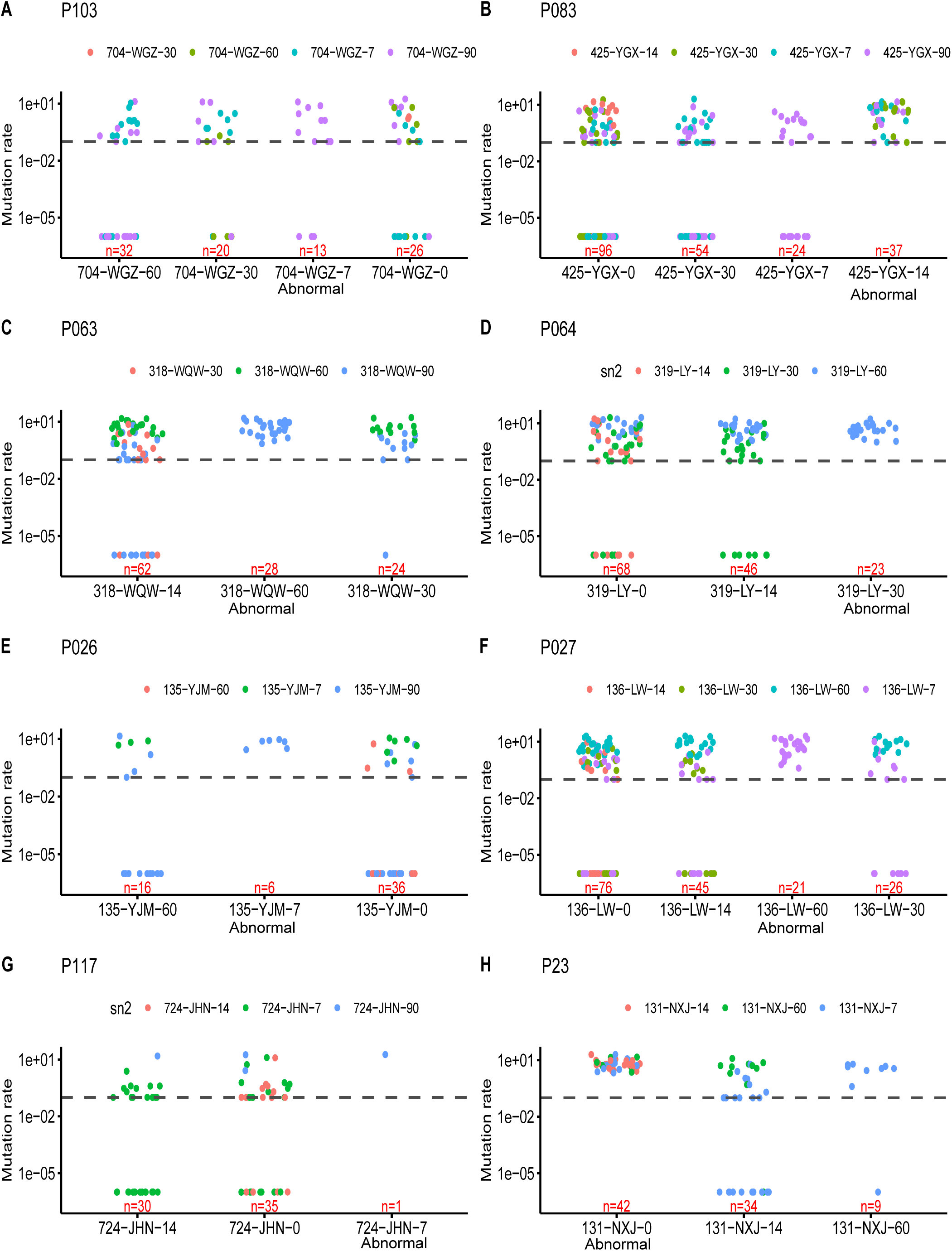
Strain-level mutation rate analysis for suspected mislabeled samples in PRJEB72385. Strain-level mutation rates (log scale) were calculated across shared (SGBs). Each panel (A–H) represents one individual with suspected mislabeled samples. The dashed horizontal line indicates the predefined strain similarity cutoff of 0.1 mutations per kilobase. Points represent mutation rates for individual shared SGBs, and sample sizes (number of shared SGBs) are shown below each comparison. Panels A show representative cases (P103) in which the abnormal samples exhibit some SGBs below the cutoff when compared with their intra-subject samples, indicating strain continuity and suggesting that these samples likely reflect true biological variation rather than mislabeling. Panels B–H show cases (P083, P063, P064, P026, P027, P117, and additional subjects) in which the abnormal samples display mutation rates predominantly above the cutoff relative to their intra-subject samples, indicating no shared strains.

**Supplementary Figure 8.**
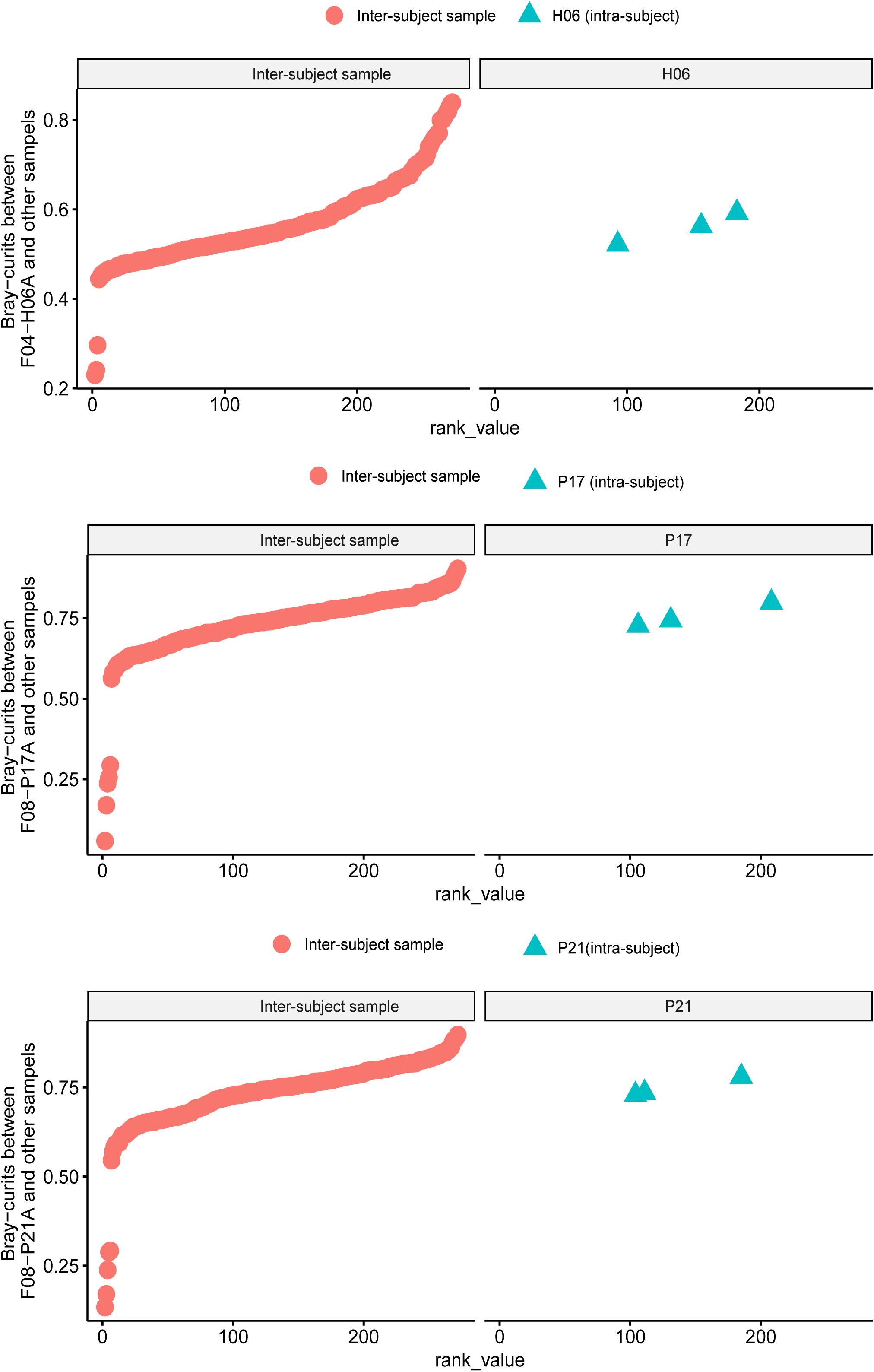
Rank-based Bray–Curtis distance analysis supporting mislabel detection in PRJCA016454. Rank distributions of Bray–Curtis distances for suspected mislabeled samples in the PRJCA016454 cohort. Each panel corresponds to one individual (H06, P17, and P21). Red circles represent inter-subject comparisons (distances between the queried sample and samples from other individuals), plotted in ascending rank order. Blue triangles represent intra-subject comparisons (distances between the queried sample and other longitudinal samples from the same individual).

**Supplementary Figure 9.**
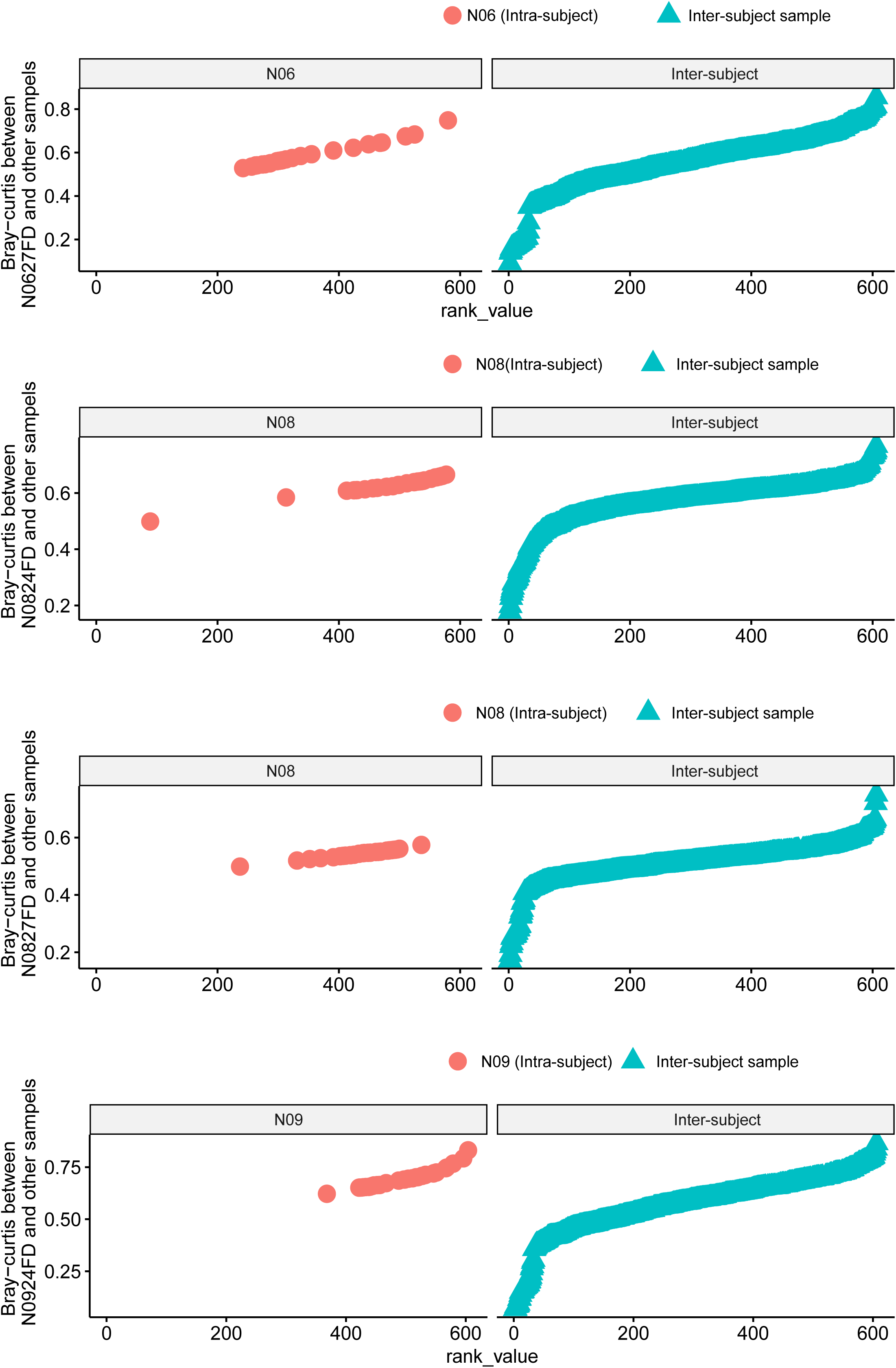
Rank-based Bray–Curtis distance analysis supporting mislabel detection in PRJCA026382 dataset. Rank distributions of Bray–Curtis distances for suspected mislabeled samples in the PRJCA026382 cohort. Each panel corresponds to one individual (N0627FD, N0824FD, N0827FD and N0924FD). Blue triangles represent inter-subject comparisons (distances between the queried sample and samples from other individuals), plotted in ascending rank order. Red cycles represent intra-subject comparisons (distances between the queried sample and other longitudinal samples from the same individual).

**Supplementary Figure 10.**
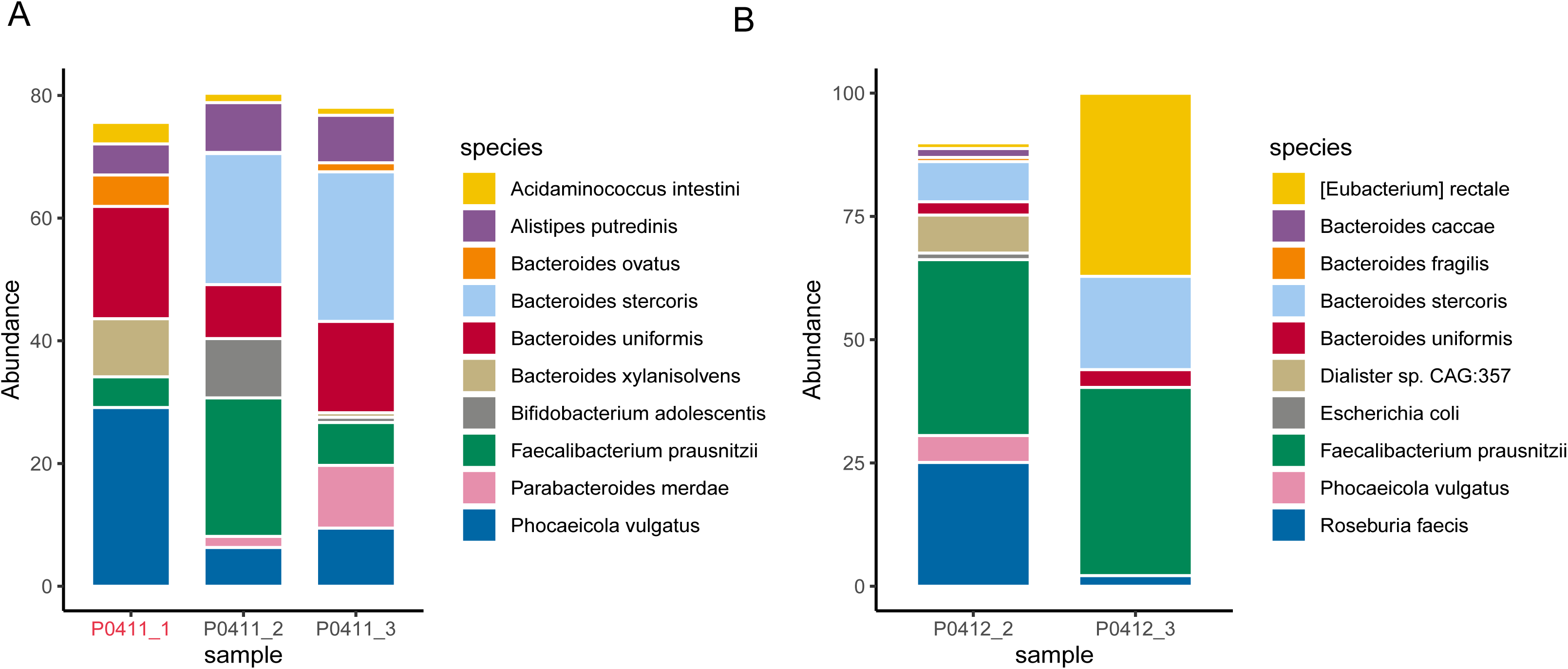
Comparison of dominant microbiome composition between suspected samples and other samples from the same subject. Stacked bar plots showing species-level relative abundance profiles for representative individuals containing suspected abnormal samples. (A) Species composition of three longitudinal samples from subject P0411. Sample P0411_1 (highlighted in red) exhibits a markedly different microbial composition compared with P0411_2 and P0411_3, including shifts in dominant taxa, indicating substantial intra-subject compositional divergence. (B) Species composition of two longitudinal samples from subject P0412. Sample P0412_3 displays a distinct community structure relative to P0412_2, characterized by major changes in dominant species.

**Supplementary Figure 11.**
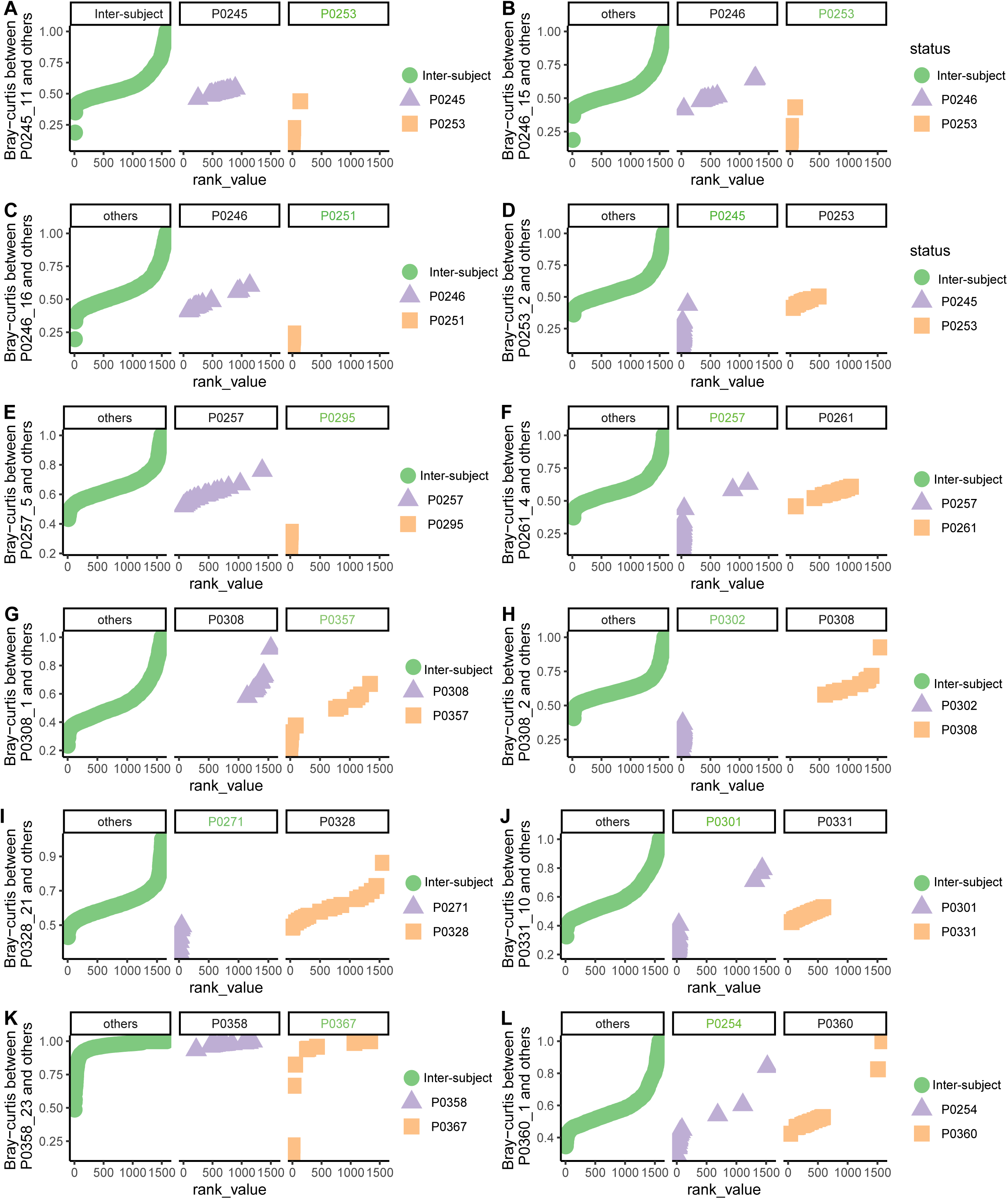
Rank-based Bray–Curtis distance analysis of suspected mislabeled samples in the HMP_2019_ibdmdb cohort. Rank distributions of Bray–Curtis distances for suspected mislabeled samples in the HMP_2019_ibdmdb cohort. Each panel corresponds to one sample. Red cycles represent inter-subject comparisons (distances between the queried sample and samples from other individuals), plotted in ascending rank order. Blue triangles or squares represent intra-subject comparisons or comparisons with the inferred true subject of origin for the mislabeled sample. The subject identified as the most likely true source is indicated in green text in the panel subtitle. Lower rank positions indicate greater similarity.

**Supplementary Figure 12.**
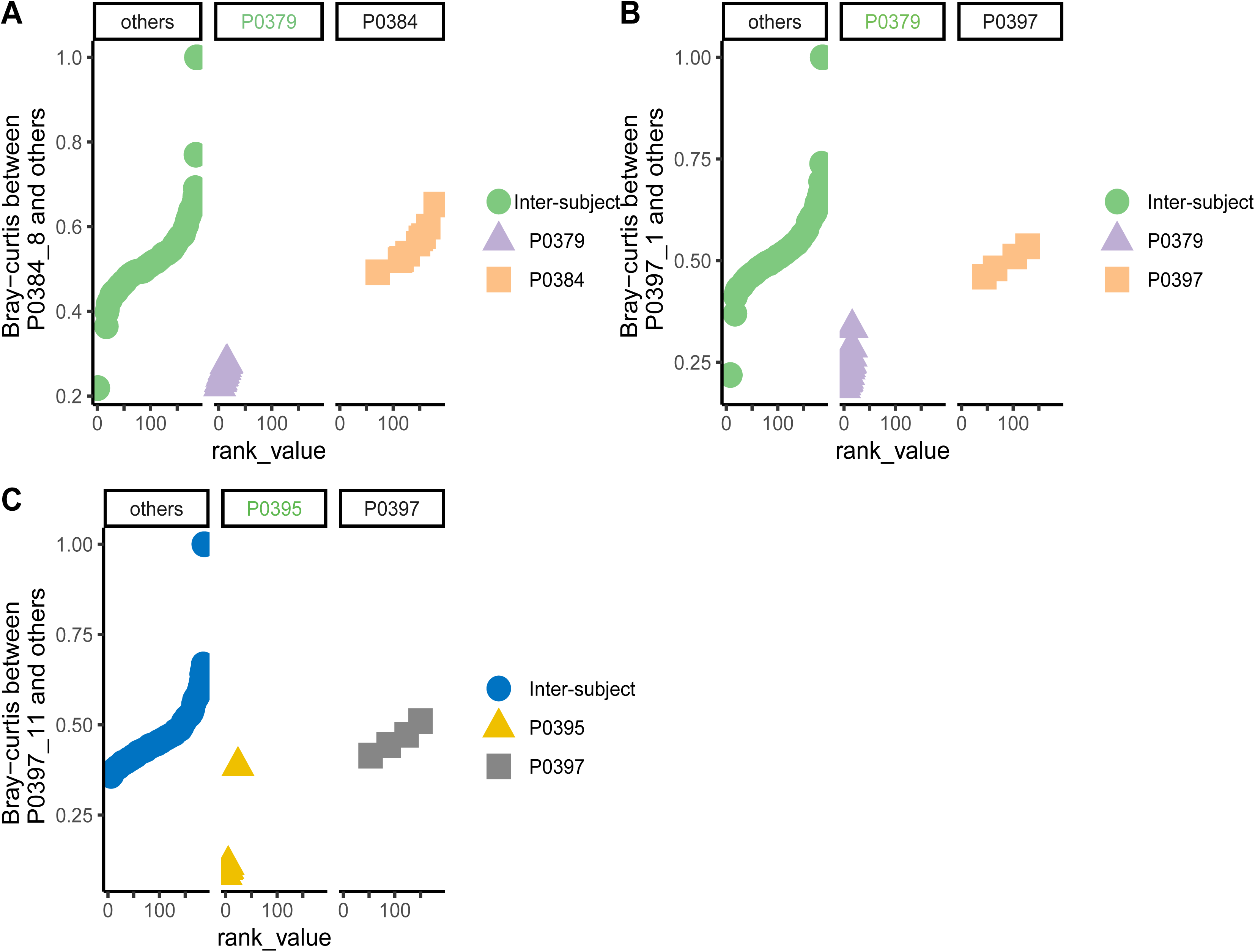
Rank-based Bray–Curtis distance analysis of suspected mislabeled samples in the HMP_2019_t2d cohort. Rank distributions of Bray–Curtis distances for suspected mislabeled samples in the HMP_2019_t2d cohort. Each panel corresponds to one sample. Cycles represent inter-subject comparisons (distances between the queried sample and samples from other individuals), plotted in ascending rank order. Triangles or squares represent intra-subject comparisons or comparisons with the inferred true subject of origin for the mislabeled sample. The subject identified as the most likely true source is indicated in green text in the panel subtitle. Lower rank positions indicate greater similarity.

**Supplementary Figure 13.**
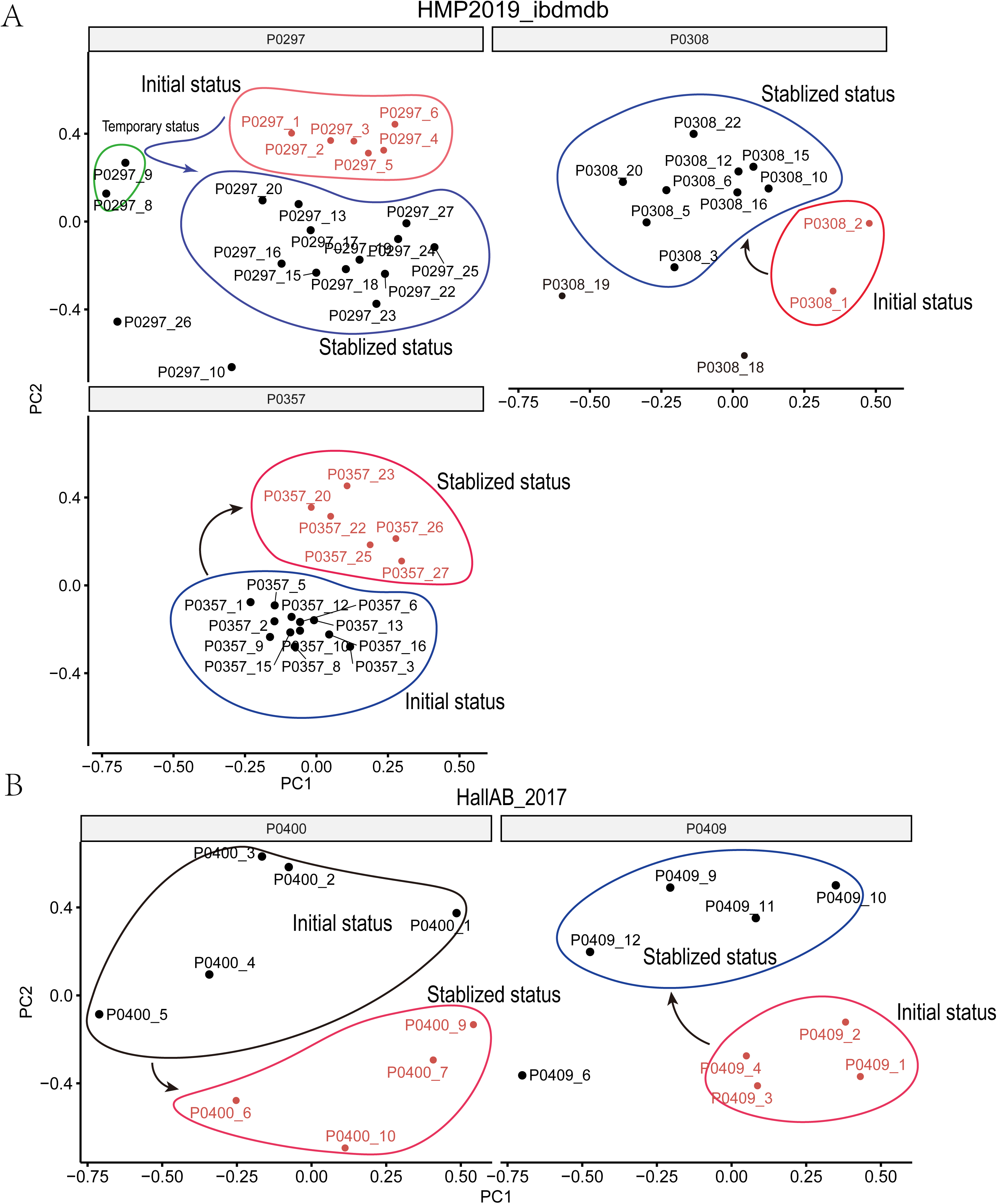
Examples of persistent and transient microbiome state transitions in longitudinal datasets. PCoA of Bray–Curtis distances illustrating temporal patterns of microbiome dynamics in individuals with consecutive abnormal samples. Each point represents a longitudinal sample, and labels indicate subject ID followed by the sampling time order (e.g., _1, _2, _3 denote sequential time points). Ellipses highlight clusters corresponding to distinct microbiome states. (A) Examples from the HMP2019_ibdmdb dataset. Individuals P0297, P0308, and P0357 illustrate two temporal patterns. In P0297, most samples cluster within a stabilized microbiome state (blue ellipse), while a small number of samples form a temporary deviation (green ellipse) before returning to the stabilized state. In P0308 and P0357, the microbiome transitions from an initial state (red ellipse) to a distinct stabilized configuration (blue ellipse), indicating persistent compositional shifts. (B) Examples from the HallAB_2017 dataset. Individuals P0400 and P0409 show similar patterns of microbiome dynamics. In P0400, the microbiome transitions from an initial state (black ellipse) to a stabilized state (red ellipse). In P0409, early samples cluster within the initial state (red ellipse), followed by stabilization into a distinct microbiome configuration (blue ellipse). Arrows indicate the temporal trajectory of microbiome changes inferred from the sampling order.

